# Potent neutralizing nanobodies resist convergent circulating variants of SARS-CoV-2 by targeting novel and conserved epitopes

**DOI:** 10.1101/2021.03.09.434592

**Authors:** Dapeng Sun, Zhe Sang, Yong Joon Kim, Yufei Xiang, Tomer Cohen, Anna K. Belford, Alexis Huet, James F. Conway, Ji Sun, Derek J. Taylor, Dina Schneidman-Duhovny, Cheng Zhang, Wei Huang, Yi Shi

## Abstract

There is an urgent need to develop effective interventions resistant to the evolving variants of SARS-CoV-2. Nanobodies (Nbs) are stable and cost-effective agents that can be delivered by novel aerosolization route to treat SARS-CoV-2 infections efficiently. However, it remains unknown if they possess broadly neutralizing activities against the prevalent circulating strains. We found that potent neutralizing Nbs are highly resistant to the convergent variants of concern that evade a large panel of neutralizing antibodies (Abs) and significantly reduce the activities of convalescent or vaccine-elicited sera. Subsequent determination of 9 high-resolution structures involving 6 potent neutralizing Nbs by cryoelectron microscopy reveals conserved and novel epitopes on virus spike inaccessible to Abs. Systematic structural comparison of neutralizing Abs and Nbs provides critical insights into how Nbs uniquely target the spike to achieve high-affinity and broadly neutralizing activity against the evolving virus. Our study will inform the rational design of novel pan-coronavirus vaccines and therapeutics.

## Introduction

Since its emergence in late 2019, the Coronavirus Disease 2019 (COVID-19) pandemic and its causative agent SARS-CoV-2 (severe acute respiratory syndrome coronavirus 2) have caused devastating consequences to global health and the economy. In response to this crisis, remarkable progress has been made by the scientific community and biopharmaceutical industry to develop innovative strategies to help curb this highly transmissible virus. In addition to vaccine development, an impressive number of neutralizing monoclonal antibodies (mAbs) have been isolated, mostly from the convalescent plasma, to facilitate a better understanding of the host immune response to SARS-CoV-2^1^. Highly potent neutralizing mAbs, both singlets and in combinations, have been approved for emergency therapeutic use with more candidates in the pipeline ^2–4^.

Likewise, antibody fragments, especially camelid V_H_H single-domain antibodies, or nanobodies have also been successfully developed for virus neutralization ^5–12^. Compared to mAbs, nanobodies (Nbs) are substantially smaller (~ 15 kDa) yet can still bind virus antigens with excellent specificity. Because of the small sizes and structural robustness, they can be easily bioengineered into multivalent forms that bind different neutralizing epitopes, thus blocking the viral mutational escape^5–8,13^. In addition, affinity-matured Nbs are generally highly soluble, stable, and can be rapidly produced in microbes such as *E.coli* or yeast cells at low costs. Stable constructs can be delivered by small aerosolized particles and inhaled for direct and highly efficient treatment of pulmonary infections ^7,8^. Most recently, this novel inhalation therapy has been successfully evaluated *in vivo* for the treatment of SARS-CoV-2 infection. At ultra-low doses, aerosolization of an ultrapotent Nb construct drastically suppresses virus infection in both upper and lower respiratory tracts and prevents viral pneumonia ^14^. Potent neutralizing Nbs represent a convenient and highly cost-effective therapeutic option to help mitigate the evolving pandemic.

Similar to other coronaviruses, the infection of SARS-CoV-2 is mediated by the spike trimeric glycoprotein (S). Each S monomer is composed of two subunits: S1 and S2. The receptor-binding domain (RBD) of S1 is critical for interacting with the host receptor angiotensin-converting enzyme 2 (ACE2). In the prefusion state, the RBD is undergoing highly dynamic switching between closed (“down”) and open (“up”) conformations on the distal tip of the spike trimer ^15–17^. In the post-fusion state, S1 shedding triggers a large conformational change of S2 to facilitate virus binding to the host membrane for infection ^18^.

The ACE2 receptor binding site (RBS) of RBD is the major target of serologic response in COVID-19 patients. As a direct adaptation against antibody pressure, however, RBS is also the primary region where a number of convergent mutations have arisen in circulating variants of SARS-CoV-2. These variants may enhance ACE2 binding leading to higher transmissibility, elude many neutralizing mAbs, including those under clinical development, and substantially reduce the neutralizing activities of convalescent and vaccine-elicited polyclonal sera ^19–21^. These adaptations led to the global emergence of convergent circulating variants of concern, including the UK strain (B.1.1.7), the South Africa strain (B.1.351), and the Brazil strain (P.1) ^22–24^. Notably, three RBS residue substitutions (K417N, E484K, and N501Y) derived from these clinical isolates have been demonstrated to drastically reduce, or abolish the binding of a large panel of neutralizing mAbs. Other examples include Y453F on the RBS (mink-human “spillover”) and non-RBS mutation N439K, which result in the immune escape from the convalescent sera ^25^. Long-term control of the pandemic will require the development of highly effective interventions that maintain neutralizing activities against the evolving strains ^22^.

Recently, we identified thousands of distinct, high-affinity antiviral Nbs that bind RBD and have determined a crystal structure of an ultrapotent one (Nb20) in complex with RBD ^8,26^. Here we assessed the impact of the convergent variants of concern and the critical RBD point mutations on the ultrapotent Nbs. Subsequent determination of 9 high-resolution structures, involving 6 Nbs bound to either S or RBD by cryo-EM provided critical insights into the antiviral mechanisms of highly potent neutralizing Nbs. Structural comparisons between neutralizing mAbs and Nbs revealed marked differences between the two antibody species.

## Results

### Potent neutralizing Nbs are highly resistant to the convergent circulating variants of SARS-CoV-2 and a super RBD variant

We performed ELISA to evaluate how 6 critical RBD mutations impact the binding of 7 highly diverse and potent neutralizing Nbs that we have previously identified ^8^. Surprisingly, the neutralizing Nbs were largely unaffected by the mutations (**Figure 1A, S1**). The only exception was E484K, which almost completely abolished the ultra-high affinity of Nbs 20 and 21. Additionally, we evaluated two circulating variants of global concern (B.1.1.7 UK and B. 1.351 SA) on Nb neutralization using a pseudotyped virus neutralization assay (**Methods**). These pseudoviruses fully recapitulate the major mutations of the natural spike variants, including deletions and point mutations (**Figure S2, Methods**). The initial SARS-CoV-2 strain (Wuhan-Hu-1) was used as a control. Consistent with the ELISA results, we found that the UK strain (B.1.1.7), possessing a critical RBD mutation N501Y, has little if any effect on all the potent neutralizing Nbs that we have evaluated (**Figure 1B, S3**). The SA strain (B.1.351), containing three RBD mutations (K417N, E484K, and N501Y), drastically reduces the efficacy of Nbs 20 and 21, but has a very marginal impact on the efficacies of other Nbs. The results contrast with recent investigations of a repertoire of neutralizing mAbs including those under clinical development, convalescent, and the vaccine-elicited polyclonal sera, which are significantly affected by at least one of these strains (**Table S1**).

**Figure 1.**
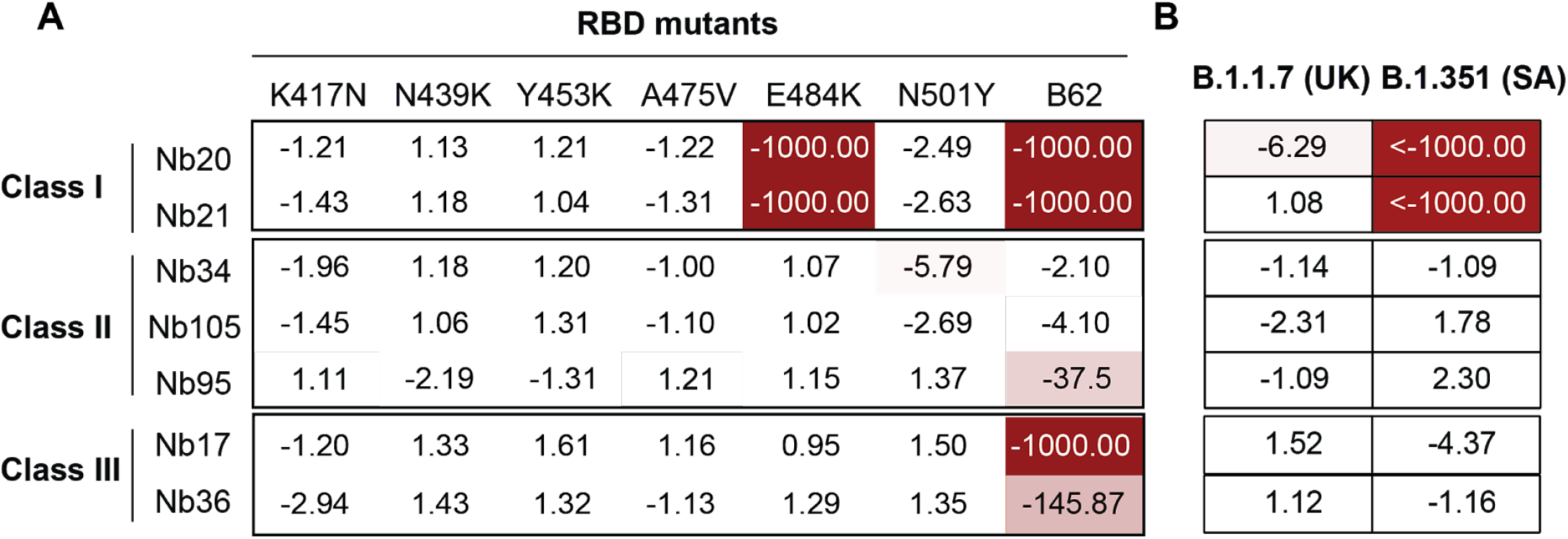
The impacts of RBD circulating variants on Nb binding. 1A: ELISA binding of the RBD mutants (a summary heatmap). Data shown as fold change of binding affinity relative to that towards RBD WT. 1B: The fold change of neutralizing potencies of the Nbs against two dominant circulating variants (UK and SA strains) compared to wild-type SARS-CoV-2 pseudovirus particles.

To assess the potential to resist future mutations, highly neutralizing Nbs were also evaluated for their binding to a potential future RBD variant (RBD62 with 9 point mutations), which was identified by *in vitro* evolution for ultrahigh-affinity ACE2 binding ^27^. While several Nbs were substantially affected by RBD62, Nbs 34 and 105 retained their high affinity against this super variant.

These striking results prompted us to further investigate the structural basis for the broadly neutralizing activities of these Nbs. Our high-resolution cryo-EM maps of Nbs in complex with either the prefusion-stabilized S ^28^ or the RBD revealed three distinct classes of neutralizing Nbs that are affected differently by the variants and provided insights into the antiviral mechanisms.

### Ultrapotent Class I Nbs and the “Achilles heel”

Class I dominates high-affinity RBD Nbs and represents some of the most potent neutralizers for SARS-CoV-2 (with the half-maximal inhibitory concentration or IC50 < 15 ng/ml). This class of Nbs is characterized by short CDR3 residues (typically 10 residues or less). Previously we have determined a crystal structure of a class I Nb (Nb20) with RBD. Nb20 can block ACE2 binding to the RBD at sub-nM concentration ^8^. Similarly, Nb21 can neutralize a clinical isolate (the Munich strain) of SARS-CoV-2 at sub-ng/ml, which is unprecedented for monomeric antibody fragments without antibody-like avidity binding ^29^. To understand the neutralization mechanism of class I Nbs better and in the context of the S trimer, we solved the structure of the most potent class I Nb (Nb21) bound to the S using cryo-EM.

Nb21 binds RBDs in both up and down conformations (**Figure 2A**). There are two major classes of the spike bound to Nb21: i) one up-RBD and two down-RBDs (resolution of 3.6 Å) and ii) two up-RBDs and one down-RBD (3.9 Å) (**Figure 2A, S4A-C**). Due to the high flexibility of RBD, we performed local refinement of one down-RBD with Nb21 to resolve the binding interface (**Figure S4C**). Nb21 binds the extended external loop region of the RBD with two β-strands. The interactions are mediated by all three CDR loops (**Figure 2B**). R31 of Nb21 forms cation-π interactions with F490 of RBD. It also forms a polar interaction network with Y104 (Nb21) and E484 (RBD). These four residues are located at the center of the Nb21:RBD interface, constituting a major site of interactions (**Figure 2C**). In addition, side chains of R97, N52, and N55 (Nb21) form hydrogen bonds with the main chain carbonyl groups of L492 and Y449 and the side chain of T470 (RBD), respectively (**Figure 2C**). The main-chain carbonyl group of A29 (Nb21) also forms a hydrogen bond with Q493 (RBD). Besides these polar interactions, F45 and L59 of Nb21 and V483 of RBD form a cluster of hydrophobic interactions, together, providing ultrahigh-affinity and selectivity for RBD binding (**Figure 2C**).

**Figure 2.**
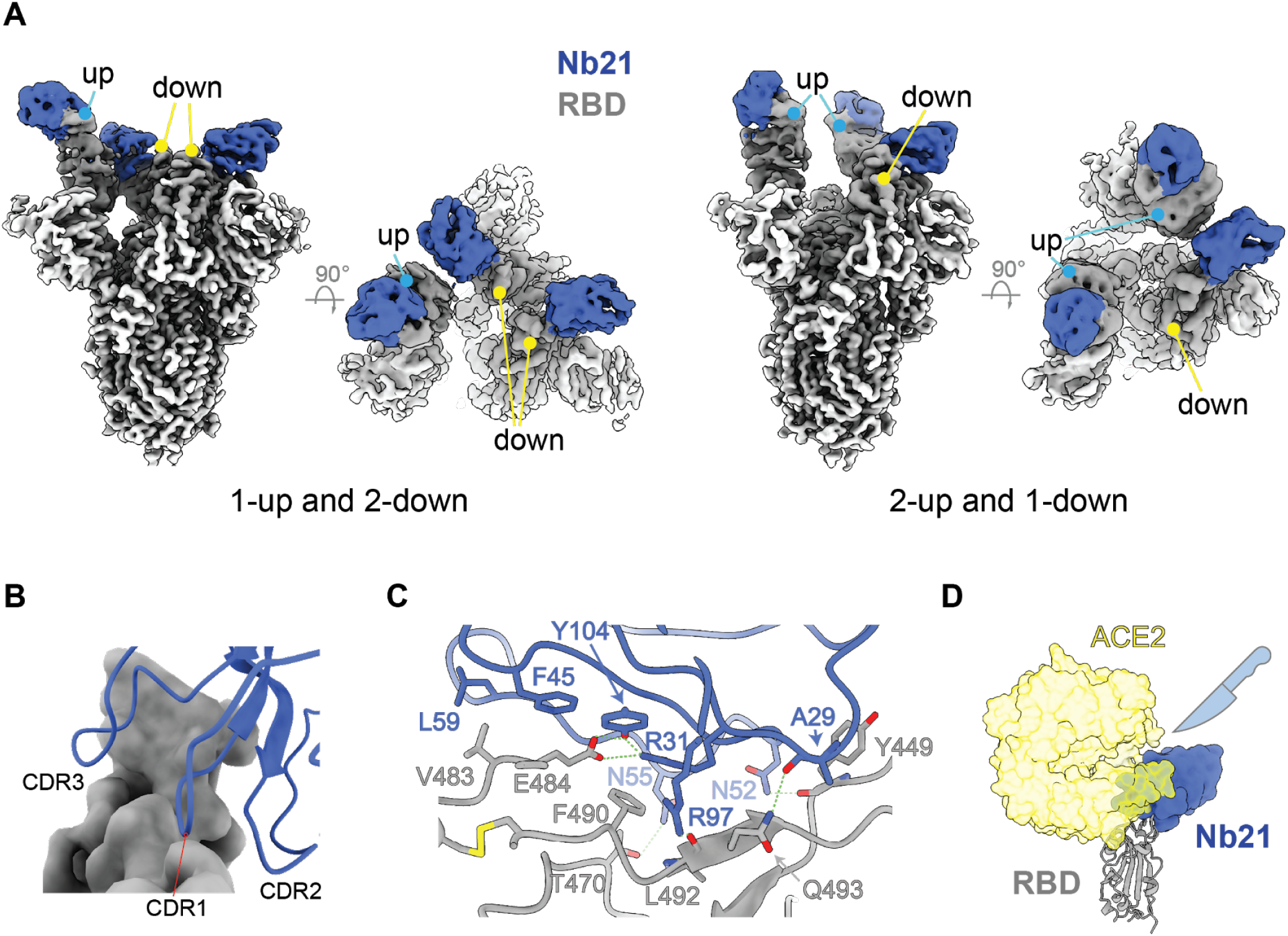
Structure of an ultrapotent class I Nb (21) 2A: Cryo-EM structure of the Nb21:S complex reveals two main RBD conformations of “1-up and 2-down” and “2-up and 1-down”. 2B: The involvement of all three CDRs of Nb21 in RBD binding. 2C: Detailed Nb21:RBD interactions. 2D: Structural overlap of hACE2 with Nb21:RBD complex.

Consistent with ELISA and pseudovirus assay results, relative binding energy calculation (**Methods**) revealed that E484 on the RBD is the “Achilles heel” of the ultrapotent Nb21 (**Figure S5A**). E484 provides the highest binding energy among all the interface residues on the RBD. In addition, it facilitates a network of adjacent residues such as F490, F489, N487, Y486, and V483 to participate in Nb21 binding. The E484K mutation can substantially destabilize the interface packing by electrostatic repulsion with R31 (CDR1), subsequently disrupting the cation-π stacking interaction between R31 and F490 (RBD). Simple charge reversal on R31(R31D) failed to recover the salt bridge and binding to the E484K mutant (**Figure 2C, S5B**). In summary, Class I Nbs bind the RBS epitopes and can potently inhibit the virus by directly blocking ACE2 binding (**Figure 2D**). Nevertheless, since the epitopes are among the least conserved regions on the spike, a critical point mutation (E484K) can dramatically reduce, or completely abolish, the ultrahigh affinity of class I Nbs (such as 20 and 21).

### Class II Nbs bind non-RBS epitopes yet still efficiently block ACE2 binding

Class II Nbs (95, 34, and 105) can potently neutralize SARS-CoV-2 below 150 ng/ml. Cryo-EM analysis of Nbs 95 and 34 with S revealed two major classes of the complexes with an overall resolution of 3.4 Å and 3.5 Å, respectively: 1) two-up-one-down RBDs (**Figure 3A**) and 2) three-up RBDs with high flexibility (**Figure S6–7**). Nb105, on the other hand, forms an elongated structure with two copies of S in all RBD-up conformations (**Figure S8**). The strong preferred orientation of this extended dimeric structure on the EM grids limits a high-resolution reconstruction to accurately define the Nb105: RBD interface. Therefore, to map the epitope, we assembled a trimeric complex of Nb21:Nb105:RBD. The resulting assembly (~60 KDa) was stable and was subsequently resolved by cryo-EM at 3.6 Å (**Figure 3B, S8**).

**Figure 3.**
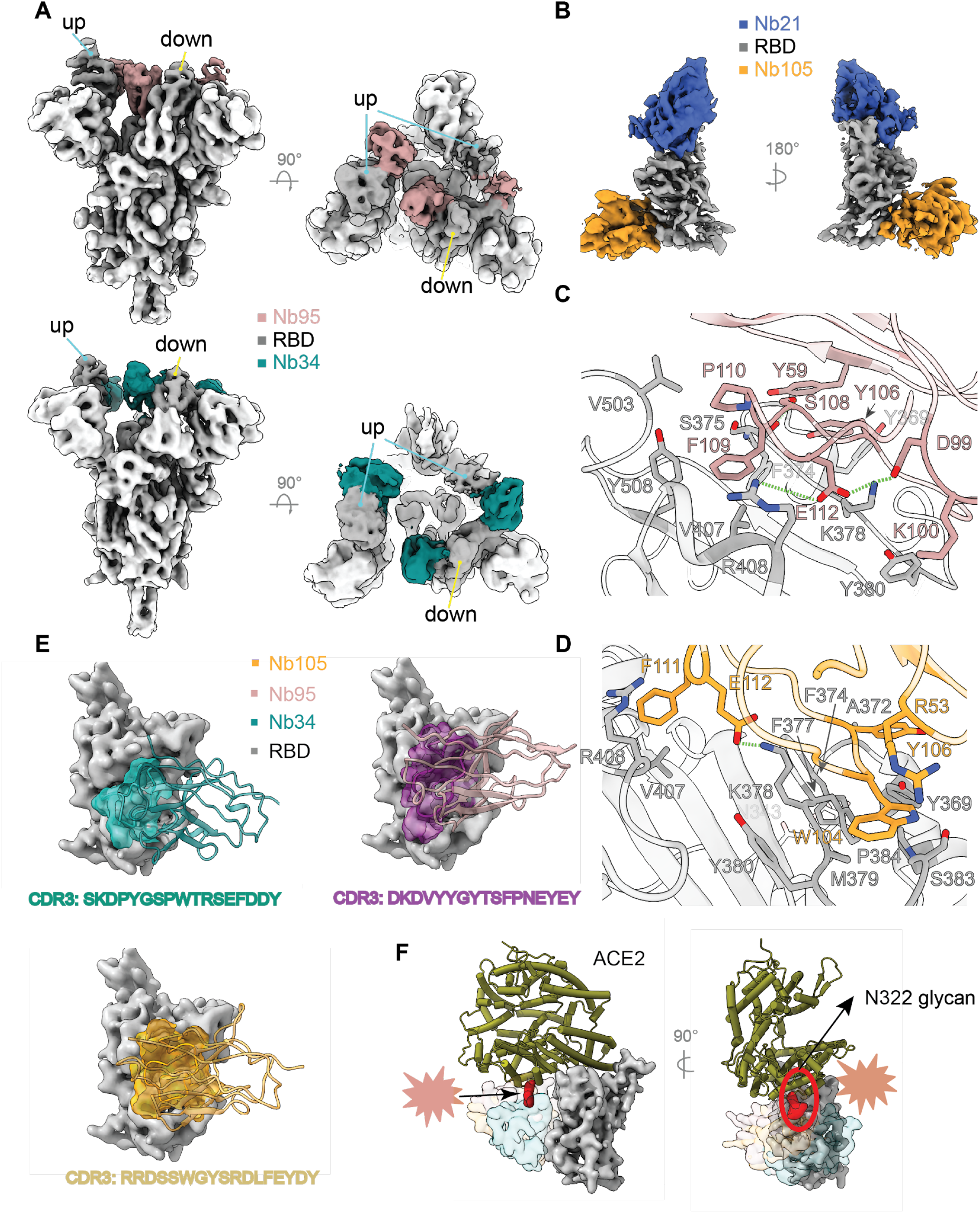
Structures of class II Nbs (95, 34, and 105) 3A: Cryo-EM structures of Nbs 95 and 34 in complex with S. 3B: Cryo-EM structure of the Nb105:Nb21:RBD complex. 3C: Nb95: RBD interactions. Residues in pink denote Nb95 for RBD binding. 3D: Nb105: RBD interactions. Residues in yellow denote Nb105 for RBD binding. 3E: Class II Nb: RBD interactions are predominantly mediated by CDR3. Nbs are represented as ribbons. The CDR3 loops are shown as surface presentations. 3F: Steric effects of class II Nbs on hACE2:RBD interactions. N322 glycosylation (ACE2) is presented in red density.

Class II Nbs bind RBD with high specificity primarily through the hydrophobic residues on a long CDR3 loop (17 or more residues). Notably, they have evolved divergent CDR3 sequences to target largely overlapping epitopes (**Figure 3E**). Major interactions of Nb95:RBD and Nb105: RBD were resolved by local refinements (**Methods**). For Nb95, the CDR3 side chains of D99, K100, and S108 form ionic or hydrogen-bonding interactions with the side chains of K378 and Y380 and the main chain carbonyl group of S375 of RBD, respectively (**Figure 3C**). R408 of RBD also forms a hydrogen bond with the main chain amine group of CDR3 E112. CDR3 residues P110 and F109 form hydrophobic interactions with V407, V503, and Y508 of RBD. Additionally, the Y106 side chain extends to interact with a hydrophobic patch formed by residues between Y369 and F374 (RBD). This interaction is further supported by a framework residue Y59, which forms hydrogen bonds with carbonyl groups of A372 and F374 of RBD (**Figure 3C**). Similar to Nb95, Nb105 recognizes RBD with CDR3 W104 and Y106 residing in two hydrophobic patches of M379-P384 and Y369-F377, respectively (**Figure 3D**). Another hydrophobic residue F111 is clamped between V407 and R408 (RBD), forming a cation-π stacking interaction. The three patches of hydrophobic interactions surround an electrostatic interaction between E112 and K378 of RBD (**Figure 3D**). Moreover, other CDRs of Nb105 help stabilize the CDR3:RBD interactions. For example, R53 (CDR2) forms a cation-π stacking on W103 (CDR3) (**Figure 3D**). While the Nb34:RBD interface was only partially resolved, it is evident that the CDR3 also plays a major role in RBD binding similar to Nb95 and Nb105 (**Figure 3E**). Superposition of three Nb: RBD structures reveals small yet different binding orientations of Nbs with respect to the RBD (**Figure 3E**).

While class II Nbs do not interact with the ACE2 directly, they can still efficiently block ACE2 binding at low nM concentrations (**Figure 4H**). Superposition of ACE2:RBD into the Nb: RBD complexes reveal that binding of Nbs 105 and 95 to the RBD overlaps with the subdomain II of ACE2 (residues 308 - 326) and the N322 glycan, wherever Nb34 can clash with the glycan (**Figure 3F**). Recent crystal structures of camelid Nbs ^5^ and synthetic constructs (sybodies)^9^ have revealed a similar mode of binding with substantially lower neutralization potency (i.e., over 100-fold) compared to affinity matured class II Nbs (95, 34, and 105). Potentially, this is due to the marked difference of these Nbs in their abilities (e.g., affinity, stability, and orientation with respect to RBD) to block ACE2 binding.

**Figure 4.**
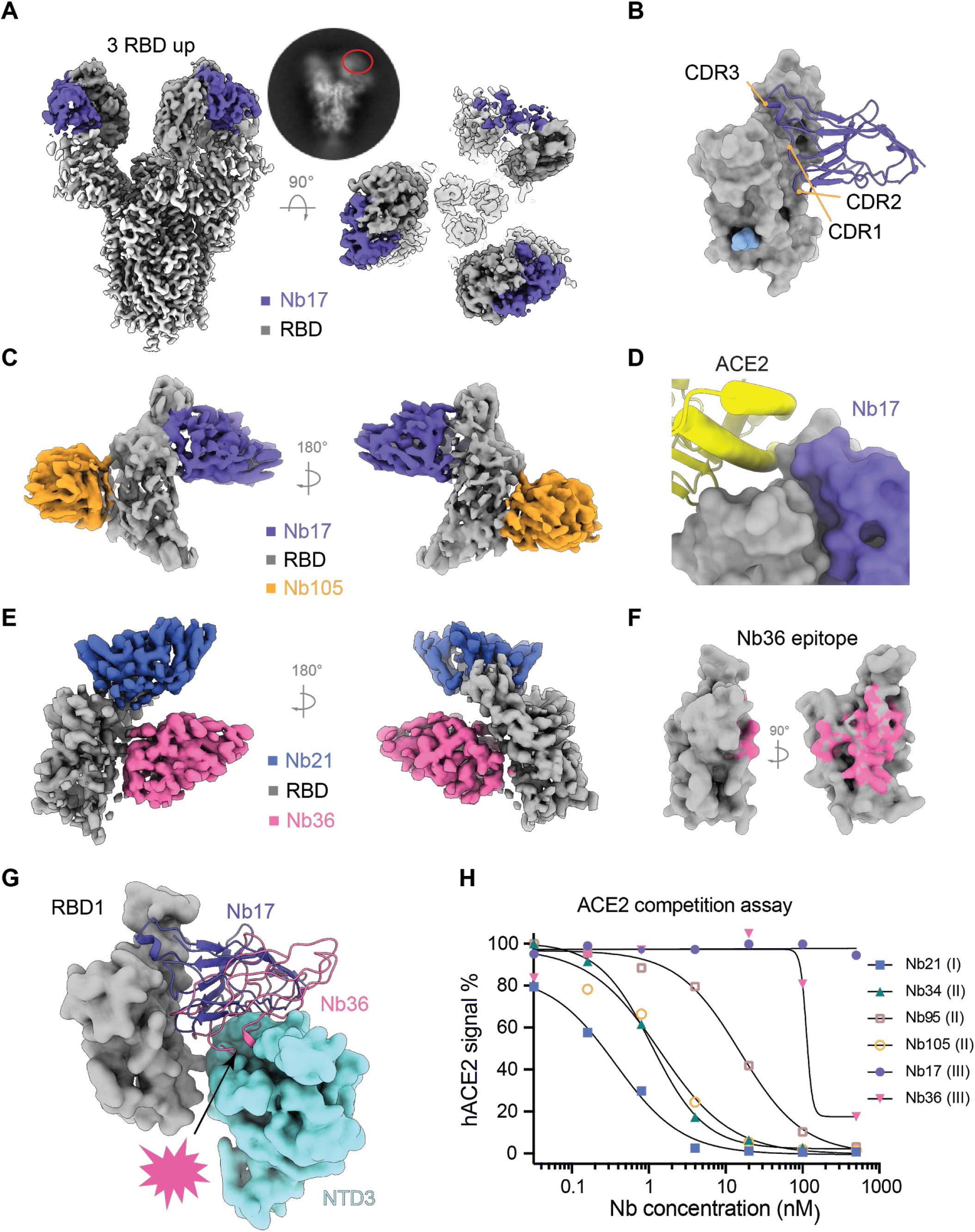
Structures of class III Nbs (17 and 36) 4A: Cryo-EM structures of Nb17 in complex with S. 4B: Nb17:RBD interactions are mediated by all three CDRs. 4C: Cryo-EM structure of the Nb17:Nb105:RBD complex. 4D: Nb17 structurally does not overlap with ACE2. 4E: Cryo-EM structure of the Nb36:Nb21:RBD complex. 4F: Epitope of Nb36 on the RBD surface. 4G: Nb17 stacks on NTD via its framework, while isolated Nb36:RBD complex indicates Nb36 would clash with neighboring NTD on S. 4H: ACE2 competition assay with the S.

### Class III Nbs utilize distinct mechanisms to neutralize the virus efficiently

#### a. Nb17 locks the spike in all RBD-up conformations

Nb17 neutralizes the virus *in vitro* at an IC_50_ of ~25 ng/ml. Notably, all three RBDs on the S trimer were in an open conformation with two having particularly strong densities (**Figure 4A, S9A-D**). Nb17 binds a conserved epitope including a segment spanning residues 345-356 and additional residues that generally do not overlap with the RBS. This epitope is localized on the opposite side of class II Nb epitopes (**Figure 4B**). Similar to Nb21, Nb17 also utilizes all three CDRs for RBD recognition. Remarkably, no bulky side chains directly involve interface packing and ultrahigh affinity RBD binding. Instead, small hydrophobic residues, such as Ala, Val, and Pro, and polar interactions are the primary contributors at the Nb17: RBD interface (**Figure S9G**). Interestingly, 3D variability analysis shows that Nb17 density stacks on the adjacent NTD prefer the open conformation of all RBDs when Nb17 is bound (**Movie S1**).

To validate our structural model, we reconstituted and analyzed the Nb17: Nb105: RBD complex to characterize the interface interactions(**Figure 4C, S9E-F**). Superposition of the Nb17: RBD complex to the ACE2:RBD complex indicates that Nb17 would not interfere with ACE2 interactions (**Figure 4D**). Structural alignment reveals that the CDR3 of Nb17 overlaps with Nb21 to compete for RBD binding (**Figure S9H**), although it remains uncertain how Nb17 neutralizes the virus. We speculate that with ultra-high affinity, Nb17 can lock S1 in a rigid and open conformation that prevents the host membrane fusion process.

Nb17 is highly resistant to all the dominant natural RBD mutations that we have tested. Compared to the Nb21:RBD interface, where E484 is buried inside the core of the interface, E484 localizes at the rim of the Nb17:RBD interface (**Figure S9H**). As such, while E484 directly contacts Nb17, the mutation (E484K) does not affect RBD binding (**Figure 1A**). The loss of binding to the super variant of RBD62 is likely caused by two point mutations (I468 and T470) (**Figure S11**).

#### a. Nb36 destabilizes the spike trimer

While Nb36:S complex is highly soluble, to our surprise, particles were not detected on the EM grids under cryogenic conditions. Therefore, to characterize Nb36:S interactions, we titrated different concentrations of Nb36 with S protein and imaged the complexes by negative stain EM. The increasing concentration of Nb36 coincided with an enhanced blurring of the particles, which compromised contrast in the electron micrographs (**Figure S10A**). This observation suggests that Nb36 can destabilize the integrity of the spike. To test this assumption, we employed thermal shift melting assays under similar conditions as those used for negative stain EM. Consistently, an increase in Nb36 concentration correlated with a decrease in protein melting temperature, indicating that Nb36 promotes instability of the S complex (**Figure S10B**). To map the epitope, we reconstituted and imaged the Nb36:Nb21:RBD complex by cryo-EM (**Figure 4E, S10C-E**). The analysis reveals that the Nb36 epitope partially overlaps with Nb17 while exhibiting no overlap with Nb21 (**Figure 4F**). The epitope covers a small segment on the non-RBS region (residues 353-360 of RBD) as well as distinct, non-RBS epitope residues that contact Nb17. Nb36 binds RBD in an orientation that is markedly different from Nb17. Superposition of the structure onto S reveals that Nb36 would have a significant steric clash with the neighboring NTD in the trimeric S complex (**Figure 4G**). Potentially, with its small size and convex epitope, Nb36 can efficiently insert between an RBD and the adjacent NTD destabilizing the highly dynamic structure of the S trimer. This destabilization mechanism is a reminiscence of mAb CR3022. However, Nb36 targets a completely different epitope from CR3022 with substantially higher neutralization potency ^30,31^.

Collectively, high-resolution cryo-EM analyses revealed both class II and III Nbs bind conserved epitopes that are incompatible with the mutational escape, thus less likely to converge into a circulating variant.

### Class III RBD Nbs belongs to a novel class of neutralizing Nbs

To investigate the RBD epitopes and Nb neutralization mechanism systematically, we analyzed all the available structures that include Nb:RBD interactions (**Figure 5A**). Epitope clustering supports the notion of three distinct classes of neutralizing Nbs. As expected, most Nbs bind class I epitopes that mainly cover RBS (**Figure 5A-B**). Class I and II epitopes are shared between Nbs and mAb (**Figure 5C-D**). In contrast, our class III Nbs are novel and unique among all the neutralizing Nbs and mAbs that have been characterized (**Figure 5E**). Class III epitopes are in close proximity to the neighboring NTD. Thus, accessing these epitopes is elusive for mAbs due to steric hindrance imposed by their large sizes. Here, with optimal orientations and substantially smaller sizes, Nbs can target this conserved region where the virus has low mutational tolerance^32^. As such, class II and III epitopes may serve as optimal targets for the development of pan-coronavirus vaccines and therapeutics.

**Figure 5.**
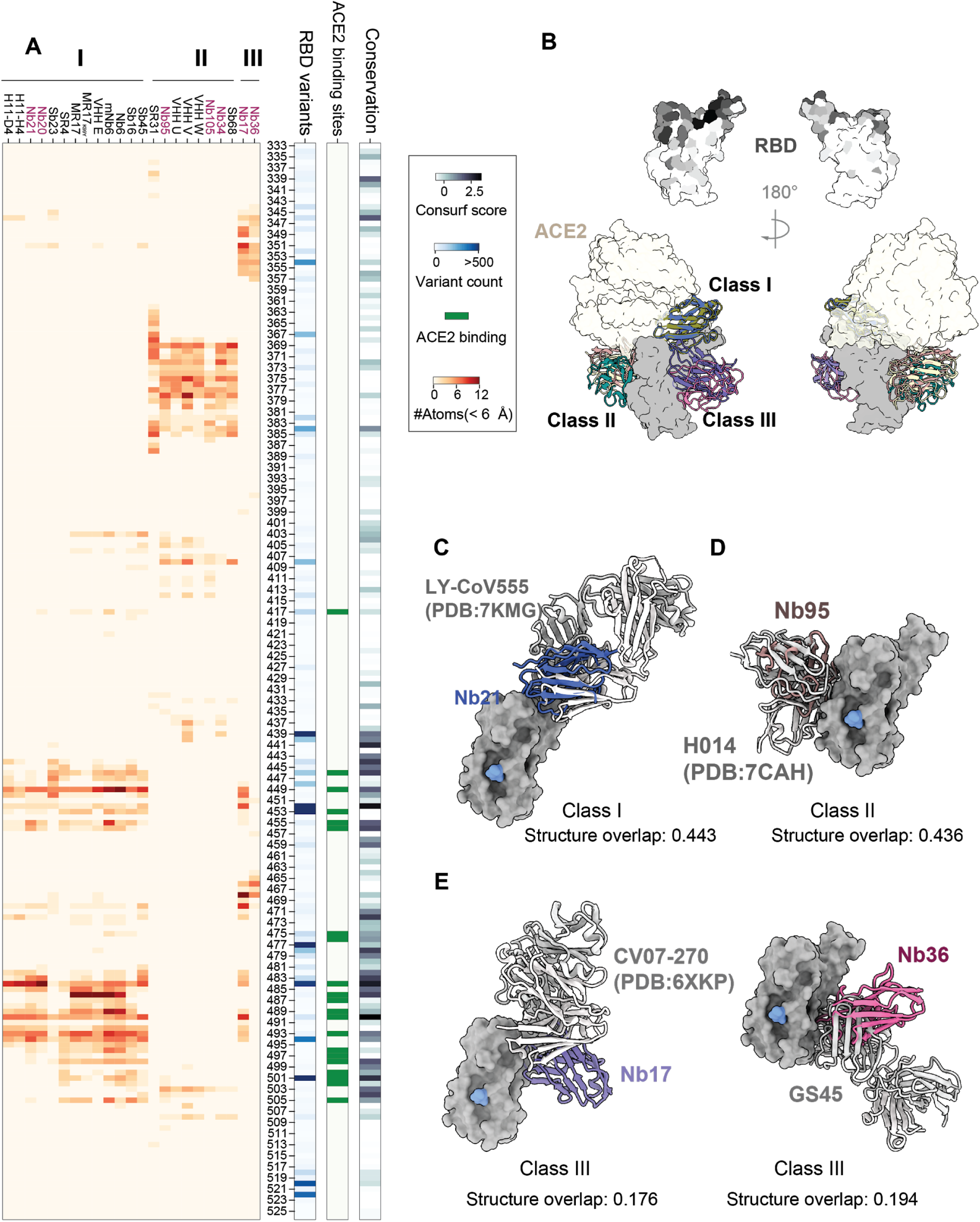
Class III Nbs bind novel and conserved neutralizing epitopes unique to Nbs. 5A: Epitope clustering analysis of RBD Nbs and correlation with RBD sequence conservation and ACE2 binding sites. 5B: Overview of three Nb classes binding to the RBD, RBD surface was colored based on conservation (ConSurf score). 5C-E: Structural comparison of different classes of Nbs with the closest mAbs for RBD binding.

### Nbs and mAbs are differently affected by mutations in the circulating variants

To understand how the unique binding modes of neutralizing Nbs are translated into their high resistance against SARS-CoV-2 mutants we compare the three Nb classes with mAbs. Buried surface area (BSA) of RBD interfacing residues from both Nb and mAb-bound structures were calculated and compared systematically (**Figure 6A-C**). The analysis reveals that the majority of mAbs (83%) use at least one of the mutated RBD residues to bind, with 60% using two or more variant residues for RBD interactions. In contrast, Nbs target these sites substantially less frequently (**Figure 6B**) with the exception of class I Nbs, which predominantly recognizes the hot spot fostered by E484 (**Figure 6D**). Other classes do not bind these variant residues directly (**Figure 6B, 6E**).

**Figure 6.**
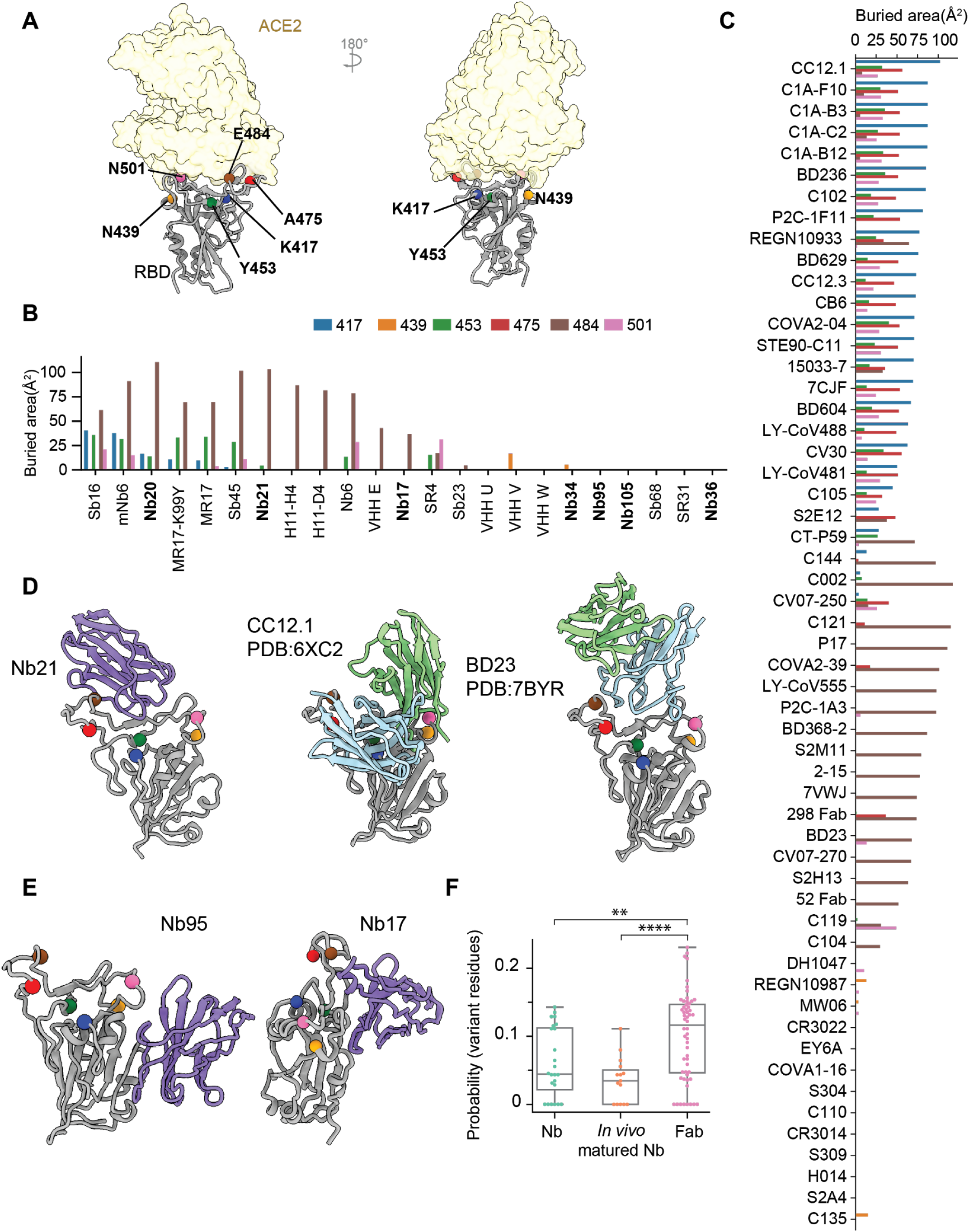
mAbs and Nbs binding to RBD are differently affected by mutations in the circulating variants. 6A: Localization of six RBD residues where major circulating variants mutate. 6B: Buried surface area of Nbs by different RBD residues. 6C: Buried surface area of Fabs by different RBD residues. 6D-E: Representative structures of different classes of Nbs with major variants residues shown as spheres. Two fab structures binding similarly to Class I Nbs were also shown on the side. 6F: The boxplot showing probability of an epitope residue to hit one of the mutations in variants.

The fact that many critical mutations localize on the RBS is intriguing (**Figure 5A**). Under selection pressures, the virus appears to have evolved a highly efficient strategy to evade host immunity by preferentially targeting this critical functional region. Specific RBS mutations (such as K417N and E484K) may help this novel zoonotic virus to optimize host adaptation (improved ACE2 binding) achieving higher transmissibility ^33^. In parallel, as RBS is the main target of serologic response, the mutations provide an effective means for the resulting variants to escape the neutralizing pressure from serum polyclonal antibodies efficiently ^27,32–34^. Since most clinical mAbs originate from convalescent plasma, it is not surprising that they are less effective to the convergent circulating variants ^19,20^. Fundamentally, this is different from neutralizing Nbs which have never been co-evolving with the virus, and therefore are less sensitive to the plasma-escaping variants. Indeed, we found that the probability of neutralizing Nb epitopes coinciding with the variant mutations was substantially lower than that of mAbs (**Figure 6F**). This is particularly the case for highly potent and *in vivo* affinity-matured Nbs, as shown in this study. Together with the functional data (**Figure 1**), our analysis provides a solid structural basis to understand how potent neutralizing Nbs can resist the convergent variants. Nbs may provide additional therapeutic benefits over mAbs for the evolving variants of SARS-CoV-2.

### Systematic comparisons of mAbs and Nbs for RBD binding

Development and structural characterization of a rich collection of neutralizing mAbs and Nbs provide an unparalleled opportunity to compare the two distinct types of antibodies for RBD binding and their respective mechanisms of action. We compiled and analyzed all the available structures including 56 distinct mAb-bound complexes and 23 Nb-bound complexes (**Table S4**). Interface residues were extracted using a 6 Å-distance cutoff to generate the residue contact profiles and epitope-paratope propensity maps. BSA and epitope curvatures were also calculated and plotted (**Methods**).

Nbs have lower BSA values than Fabs (μ_Nb_=779 Å^2^ v.s. μ_Fab_=862 Å^2^,p=0.055) (**Figure 7A**). In addition, the distribution of BSA for Nbs is distinct from Fabs and is substantially narrower (σ_Nb_ = 151 Å^2^,σ_Fab_ = 210 Å^2^) ^35^ (**Figure 7A**). Despite the smaller size, Nbs have evolved multiple strategies for high-affinity RBD binding. They exploit surface residues (especially using CDR3 loops) significantly more efficiently than Fabs for RBD engagement (**Figure 7B-C**). Nbs also have higher BSA per-interface residue (**Figure 7B**). The involvement of the framework (FR) regions in RBD binding, particularly FR2, is also evident probably due to the absence of light chain pairing (**Figure 7C**). Interestingly, we found that compared to *in vivo* affinity-matured RBD Nbs, *in vitro* selected Nbs tend to use highly conserved FR sequences more extensively for interactions. More dominant involvement of conserved FR likely suggests that *in vitro* selected Nbs may interact less specifically to RBD (**Figure 7E, 7G**). Compared to Fabs, Nbs bind more concave surfaces (**Methods**) to tighten the interactions (**Figure 7D, 7F**). Finally, neutralizing Nbs employ electrostatic interactions more extensively while both types of antibodies predominantly use hydrophobic interactions to achieve high specificity (**Figure S12–13**).

**Figure 7.**
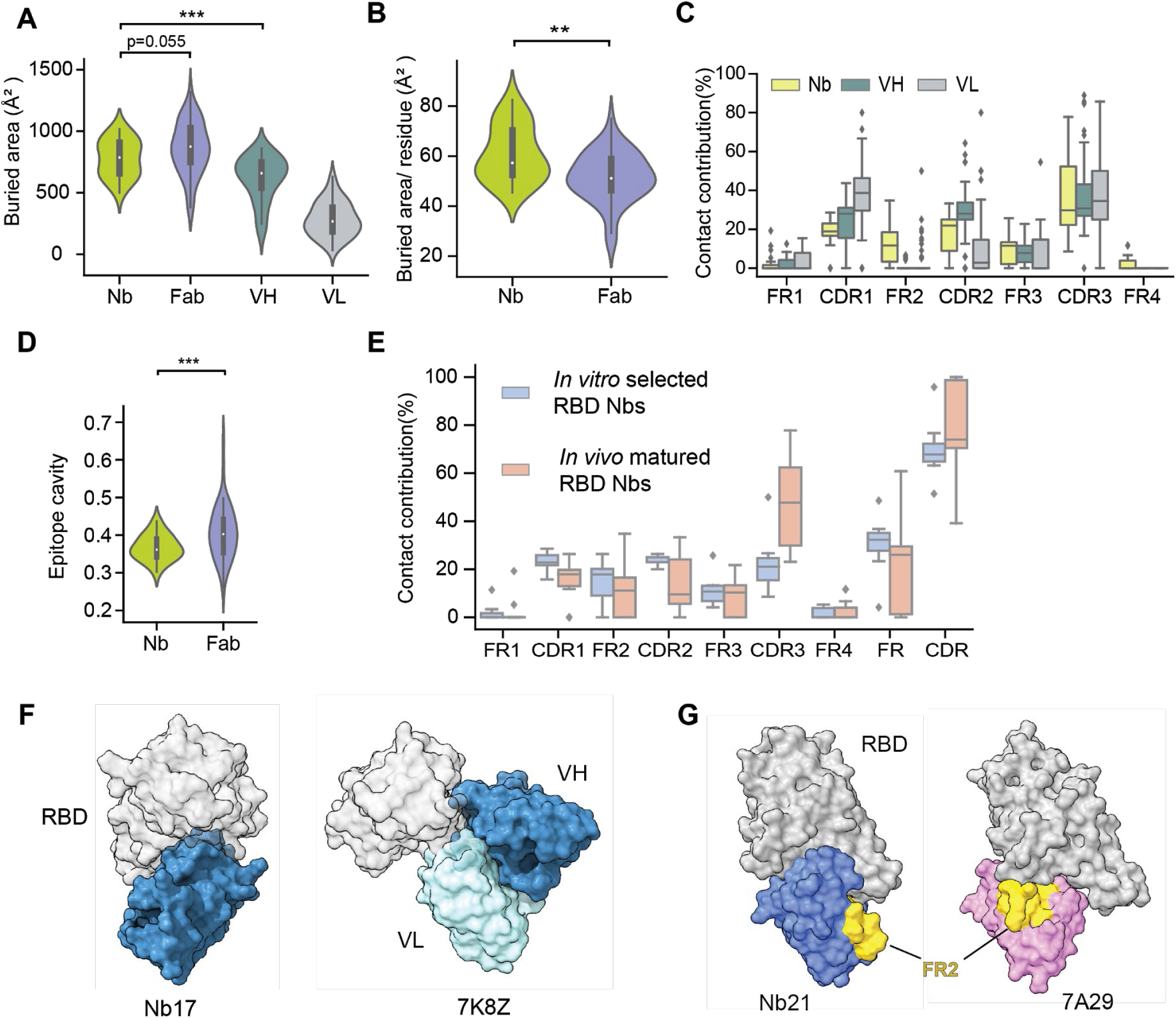
Comparisons of RBD neutralizing Nbs and mAbs. 7A: Buried surface areas of RBD: Nb and RBD: Fab complexes. VH: heavy chain. VL: light chain. 7B: Buried surface areas per-interface residue for Nbs and Fabs. 7C: The contact contribution of CDRs and FRs of Nbs and Fabs in RBD binding (using a 6 Å cutoff). Contact contribution % was calculated as # of contacting residues on CDR or FR region/total # of contacting residues. 7D: Quantification of interface cavity. Y-axis is the curvature value. 7E: Comparison of contributions from CDRs and FRs for RBD binding between *in vivo* matured Nbs and *in vitro* selected Nbs. 7F: Representative structures showing different binding modes (epitope curvature) of Nb17 and a Fab. Nbs target concave RBD surfaces to achieve high-affinity binding. 7G: Representative structures showing the direct involvement of FR2 from an *in vitro* selected Nb (PDB# 7A29) for RBD interaction.

## Discussion

By March 2021, over 116 million people had been infected by SARS-CoV-2. While the virus continues to rage with hundreds of thousands of daily new infections, the prospect of curbing it rests on the development of effective interventions including vaccines and therapeutics that are broadly neutralizing to resist both the current and future circulating variants. Highly selective neutralizing Nbs and the multivalent Nb forms represent some of the most potent antiviral agents ^5–8,13^. They are characterized by marked stability and solubility to facilitate robust and low-cost manufacturing and storage, which are essential during the pandemic. Critically, the stable constructs can be delivered by novel aerosolization route to treat virus pulmonary infection with high efficacy ^14^.

Compared to neutralizing mAbs that have been systematically characterized, structures and mechanisms for highly potent neutralizing Nbs had yet to be investigated. In this work, by using single-particle cryo-EM, we have solved a repertoire of such Nbs in complex with the spike or RBD. Neutralizing Nbs can be grouped into three epitope classes. Class I Nbs are the most abundant neutralizing Nbs and bind RBS, which is one of the least conserved regions on the spike. They can bind both up and down RBD conformations on the spike (**Figure 2**). While class I Nbs can neutralize the virus at extremely low concentrations (e.g., sub-ng/ml), their RBD binding can be abolished by a single point mutation E484K. Class II and III bind conserved epitopes that are not accessible in the closed RBD conformation. Class II Nbs can bind 2-up-1-down RBD conformations. They bind non-RBS epitopes yet can still sterically block ACE2 binding (**Figure 3**). Class III Nbs bind novel and conserved epitopes that have not been previously discovered (**Figure 4**). The cryptic RBD epitopes are in the vicinity of NTD that are highly challenging to access by large mAbs. While Nb36 can potentially neutralize the virus by destabilizing the spike, it remains unclear how the highly potent class III neutralizer Nb17 functions. Presumably, Nb17 can promote the shedding of S1 to inhibit virus-host entry, with ultra-high affinity. Nevertheless, the unique RBD binding mode of Nb17 and its ease of bioengineering may facilitate the development of robust biosensors to detect specific conformations of the spike. Currently, this process can only be inferred computationally ^36–39^ or by single-molecule techniques which require extensive bioengineering to label the spike with fluorescence dyes ^17^. Finally, since Nb17 binds extremely tightly to the RBD-up conformations, potentially, it could be used as a novel Nb “adjuvant” for vaccines to facilitate exposure of conserved yet cryptic RBD epitopes that could help elicit broadly neutralizing activities insensitive to the evolving variants.

The convergent variants emerged from persistent serologic pressure *in vivo* ^40–42^, and current interventions were developed against the initial SARS-CoV-2 strain that emerged in 2019. It is therefore not surprising that the prevalent variants can evade most clinical antibodies and weaken the polyclonal antibody activities from the vaccine-elicited sera. Compared to mAbs, however, Nbs originate from camelids or are identified by *in vitro* affinity maturation. With small sizes as well as distinct structural and physicochemical properties, Nbs can target conserved and unique epitopes on the highly dynamic spike structure that are likely fundamentally inaccessible by mAbs. Our systematic analysis including both high-resolution structural studies and functional data suggests that potent neutralizing Nbs are highly resistant to the convergent circulating variants, and possibly future variants that may occur under vaccine-elicited herd immunity (**Figure 1**). Our structure-function investigations provide a framework to map neutralizing epitopes systematically and to understand the mechanisms by which Nbs efficiently inhibit the virus and its variants (**Figure 6D-F**).

The novel structural information presented here may also help rational design of “pan-coronavirus” therapies and vaccines. The breadth and potency of class II and III Nbs could be further improved by *in vitro* affinity maturation using phage or yeast display methods. Guided by the structures, it is now possible to generate multivalent constructs by fusing specific class II and III Nbs with a proper length(s) of flexible linkers. Such constructs may enable avidity binding to substantially improve the neutralization potency while locking the virus on the highly conserved sites with a very low tolerance for future mutations. Combining these multivalent constructs into novel cocktails may provide the ultimate protection from the mutational escape ^8^. Future studies will be needed to test these ideas and critically, evaluate the *in vivo* efficacies of these novel inhalable constructs in both preclinical and clinical settings ^14^.

---

**Figure S1:**
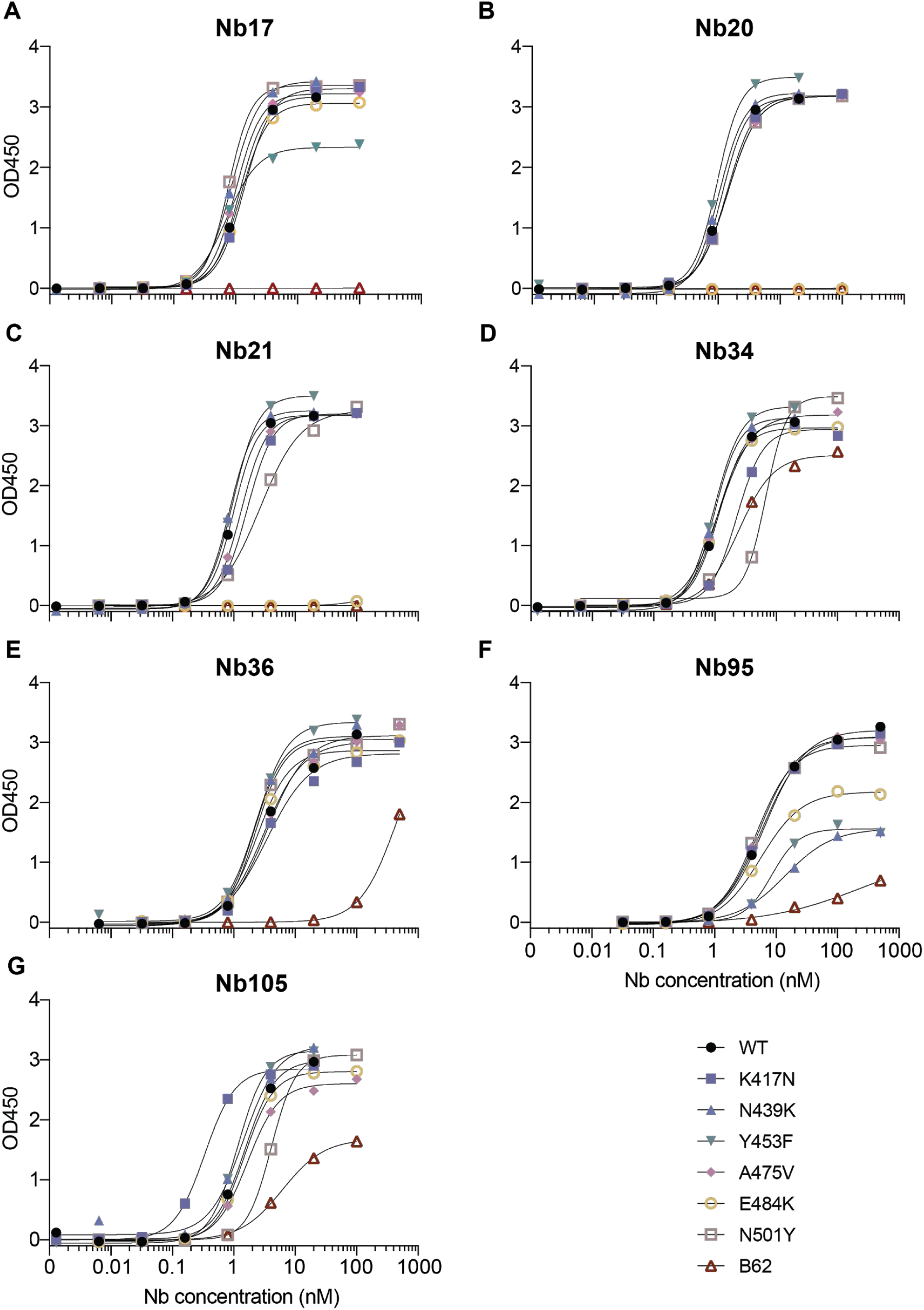
ELISA curves of Nbs for RBD mutant binding (related to Figure 1).

**Figure S2:**
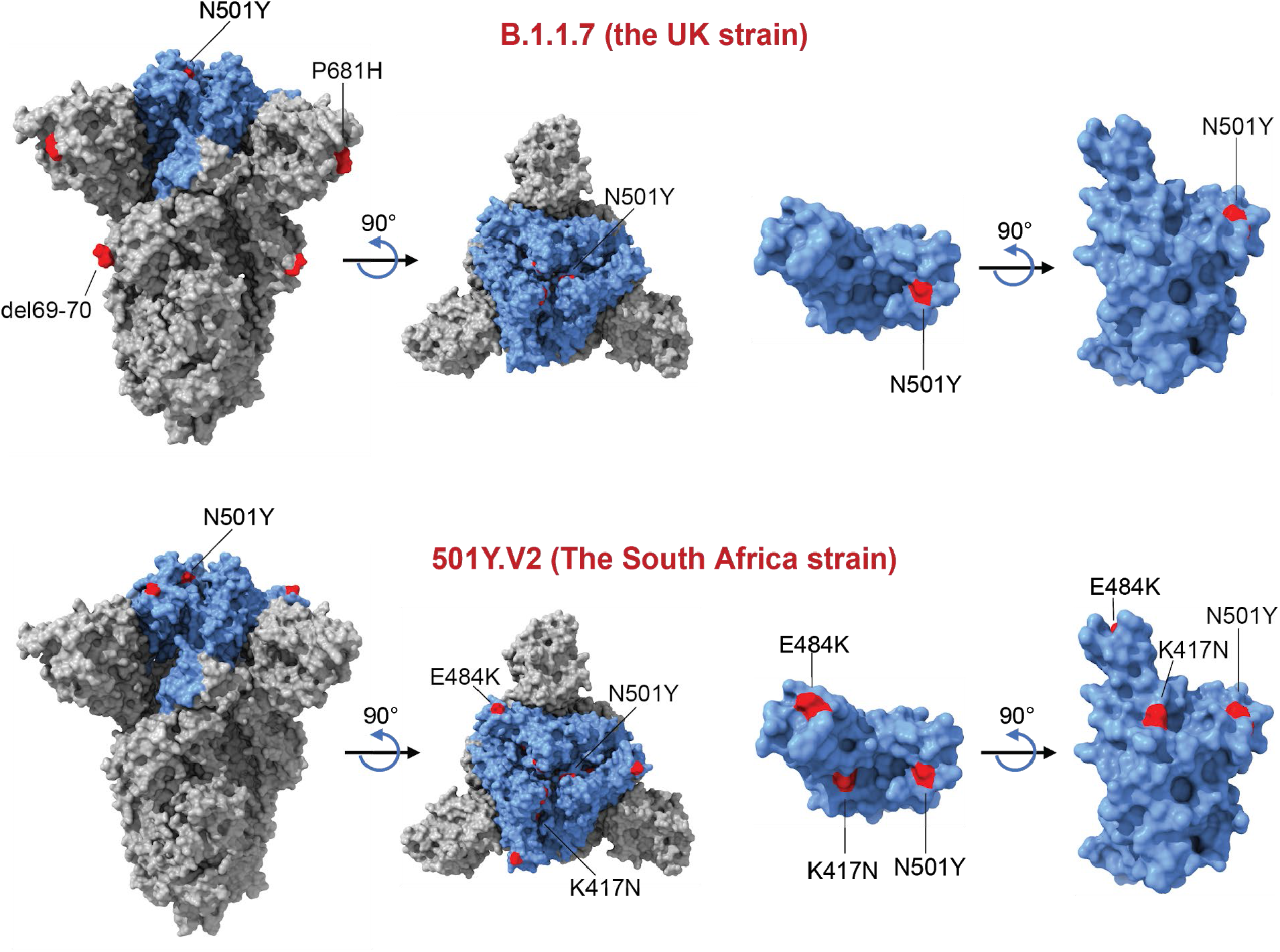
Structure representations of SARS-CoV-2 spike trimer glycoprotein and mutations for two prevalent circulating strains (related to Figures 1 and 5).

**Figure S3:**
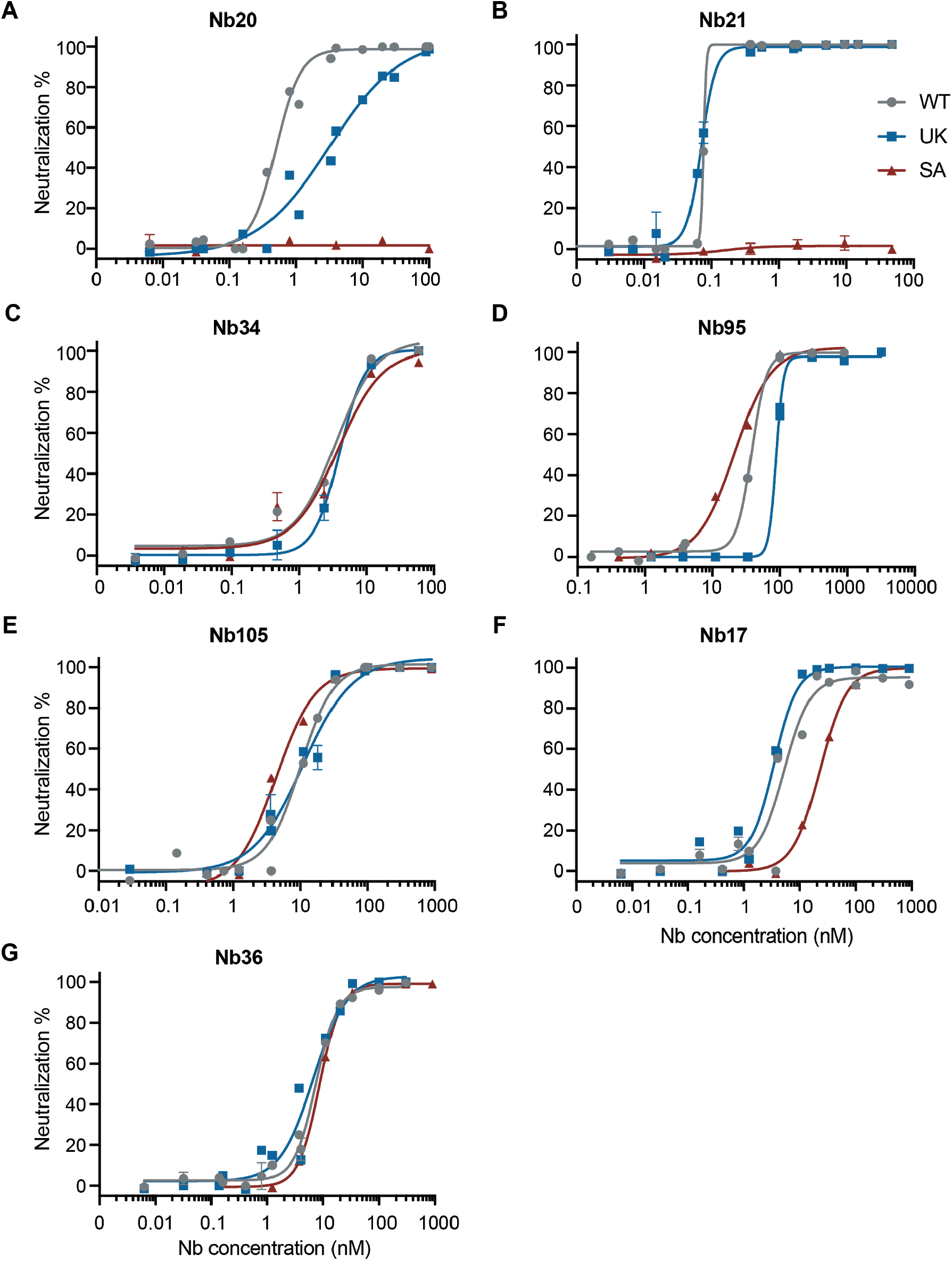
Pseudovirus assay results for individual Nbs (related to Figure 1).

**Figure S4.**
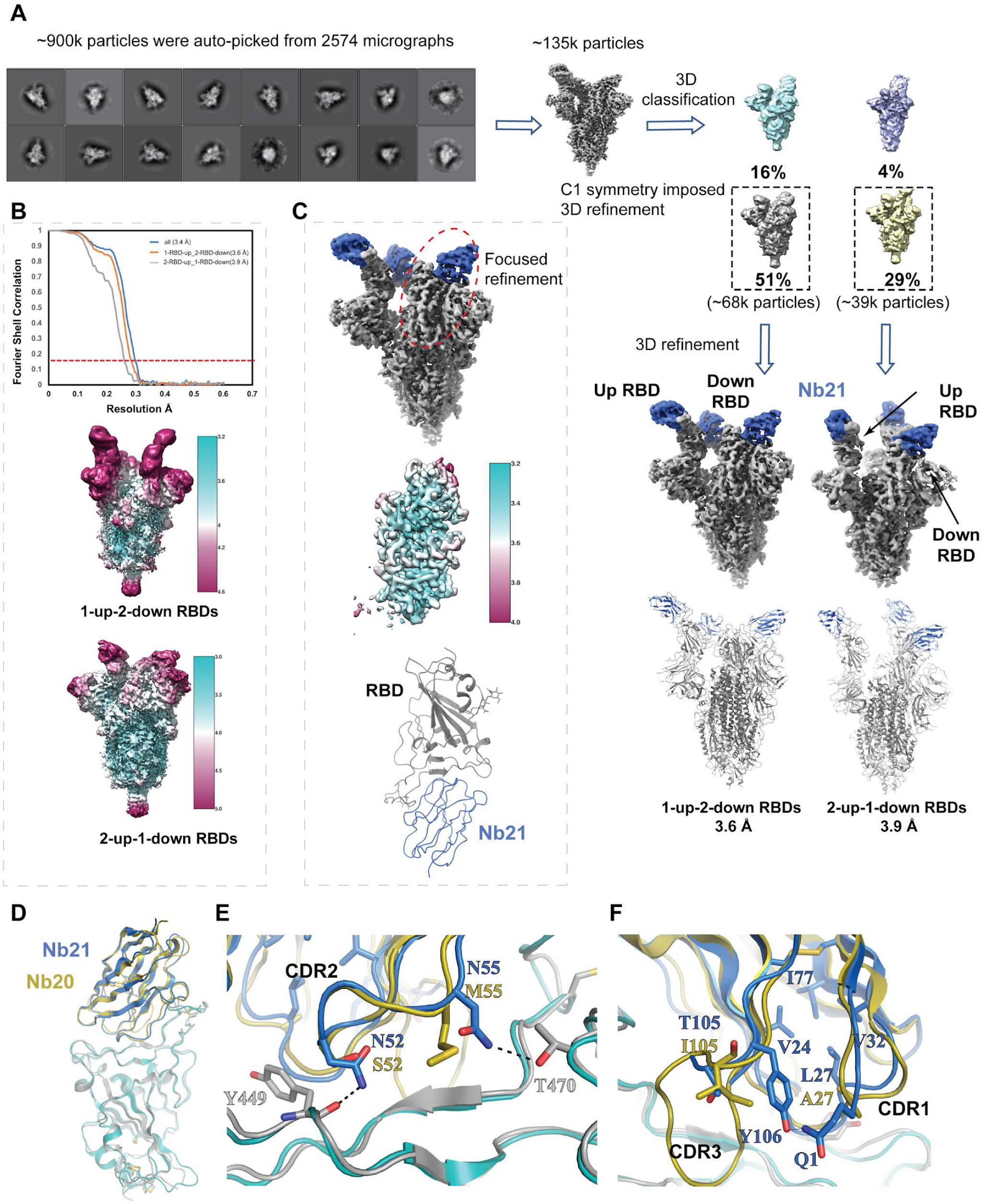
Cryo-EM Structure determination of S with Nb21, local refinement of RBD with Nb21 and comparison of Nb21 with Nb20 (related to Figure 2). (A) Cryo-EM Data Processing workflow showing the strategies and particle cohort sizes used to generate the maps discussed in this work. ~900K particles were picked based on the 2D class averages of S with Nb21 for 3D classification. Two major classes with the largest proportions were further refined. One class refined to 3.6 Å corresponds to S with 1-up-2-down RBDs and the other class refined to 3.9 Å corresponds to S with 2-up-1-down RBDs. S is colored in dark gray. Nb21 is colored in blue. (B) Fourier Shell Correlation and local resolution estimations for S and NB21 complexes. The red line represents FSC = 0.143. (C) Focused Refinement of one down RBD with Nb21. (D-F) Structural comparison of RBD with Nb21 and with Nb20. Nb21 is colored blue while Nb20 is colored yellow. RBD is colored dark gray and cyan in the structures with Nb21 and Nb20, respectively. Nb21 differs from Nb20 by four residues (all on CDRs). Its RBD binding is very similar to that of Nb20. The two structures can be well aligned with a root mean square deviation (RMSD) of 1.8 Å (all atoms) (Figure S4D). Here, S52 and M55 on CDR2 in Nb20, are replaced by N52 and N55 in Nb21, which form additional polar interactions with the RBD (Figure S4E). A27 (CDR1) and I105 (CDR3) of Nb20 are replaced by L27 and T105 in Nb21 (Figure S4F). While the two residues do not bind RBD directly, the side chain of L27 is buried inside Nb21 to form additional hydrophobic interactions with V24, V32, and I77. The small short side chain of T105 allows the neighboring residue Y106 to point towards the first N-terminal residue Q1 to form a hydrogen bond. This interaction, which is missing in the structure of Nb20:RBD as it is impeded by the presence of the large side chain of I105 in the analogous position of Nb20 I105. These additional interactions may help stabilize CDR1 and CDR3 loops to strengthen Nb21:RBD interactions.

**Figure S5.**
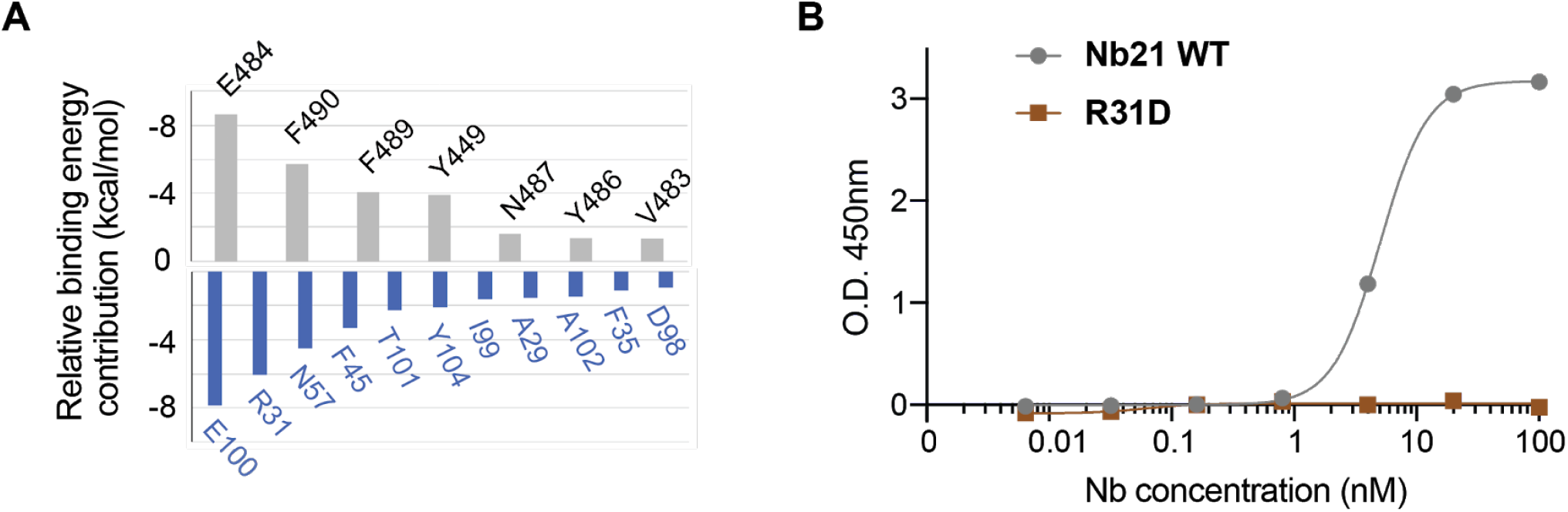
Assessment of the RBD:Nb21 interactions using both computational binding energy calculation and experimental mutagenesis (related to Figure 2). (A) Decomposition of relative binding free energy contribution from individual residues of RBD (top, gray) and Nb21 (bottom, blue) for these more than −1 kcal/mol. (B) ELISA assay showing Nb21 point mutant R31D fails to bind RBD.

**Figure S6.**
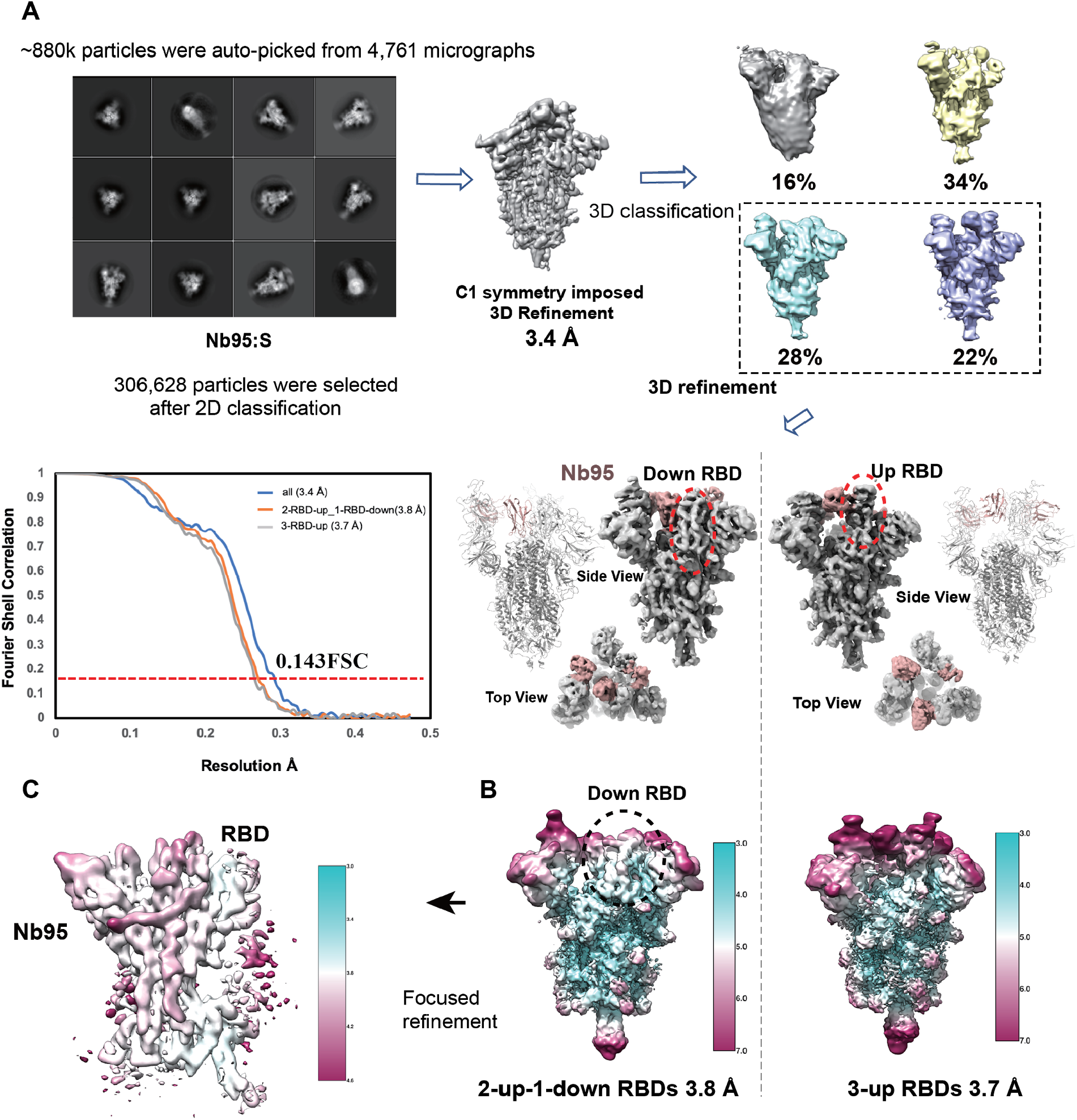
Cryo-EM Structure determination of S with Nb95 and focused refinement (related to Figure 3). (A) ~880K particles were picked based on the 2D class averages of S with Nb95 for 3D classification. Two major classes with the largest proportions were further refined. One class refined to 3.8 Å corresponds to S with 2-up-1-down RBDs and the other class refined to 3.7 Å corresponds to S with 3-up RBDs. S is colored in dark gray. Nb95 is colored in teal. FSC estimations for these two complexes are shown at the lower-left corner. The red line represents FSC=0.143. (B) Local resolution distribution for the two S and Nb95 complexes. (C) Focused Refinement of one down RBD with NB95. The down RBD showed better density compared to up RBDs.

**Figure S7.**
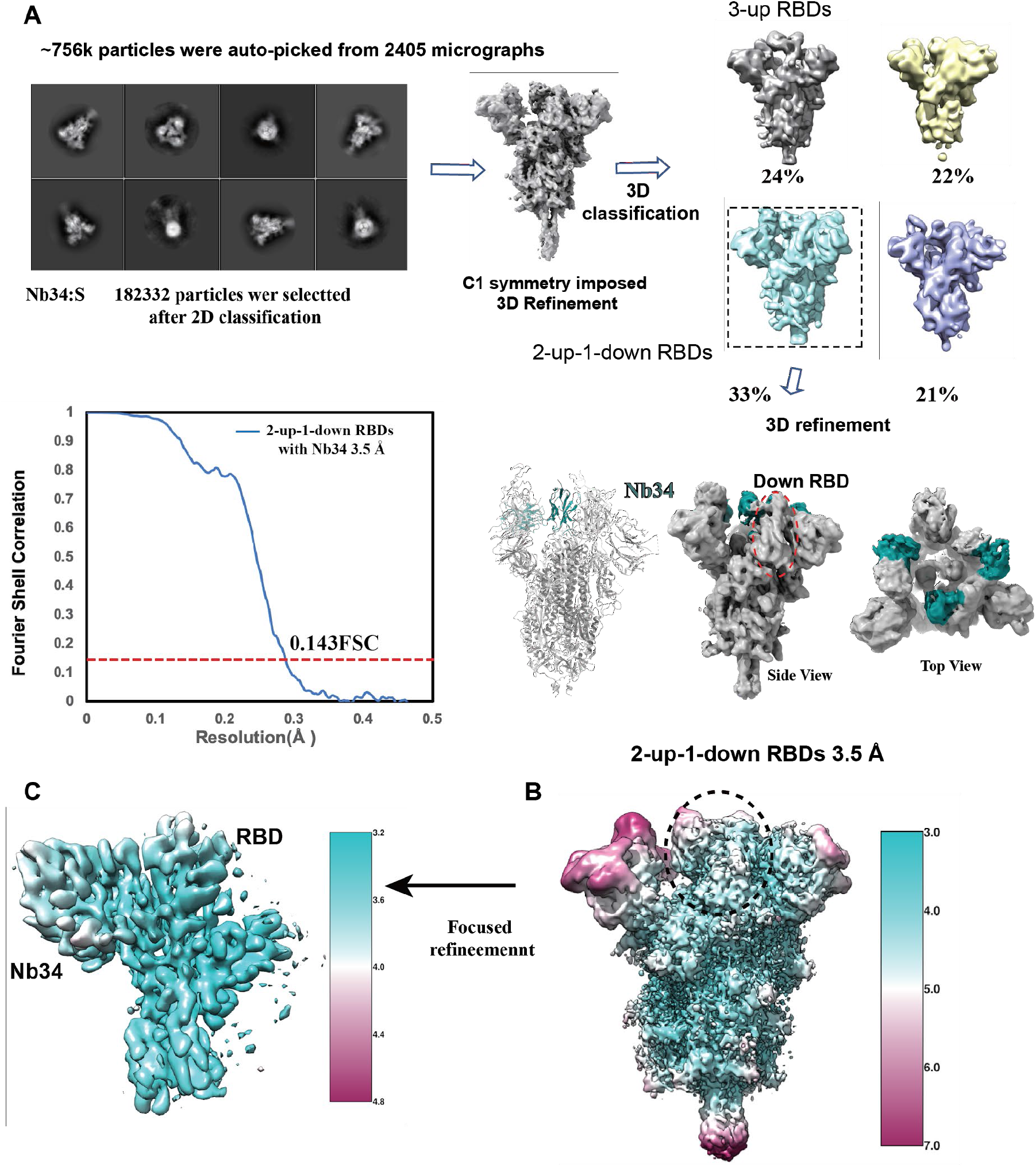
Cryo-EM Structure determination of S with Nb34 and focused refinement (related to Figure 3). (A) ~756K particles were picked based on the 2D class averages of S with Nb34 for 3D classification. Two major classes were observed with clear features of 3-up RBDs and 2-up-1-down RBDs. We focused on the 2-up-1-down class for 3D refinement and obtained a structure with a global resolution of 3.5 Å corresponding to 0.143FSC shown at the lower-left panel. (B) Local resolution distribution for the S and Nb34 complex. (C) Focused Refinement of one down RBD with NB34. The down RBD showed better density compared to up RBDs.

**Figure S8.**
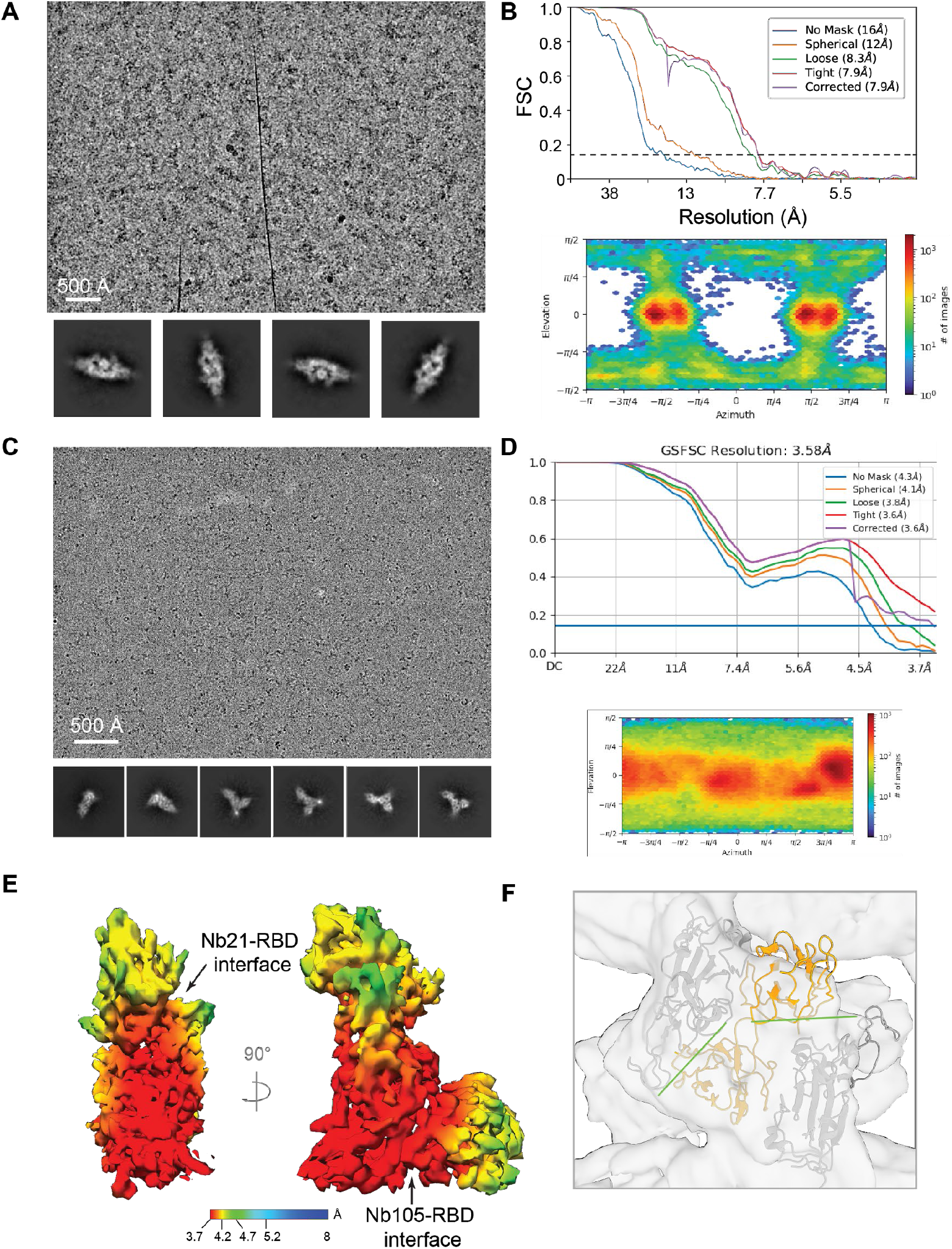
Cryo-EM analysis of Nb105:S and Nb105:RBD: Nb21 complexes (related to Figure 3). (A) Representative micrograph and 2D class averages of Nb105:S complex. (B) Gold-standard Fourier shell correlation (FSC) and Euler angular distribution. (C) Representative micrograph and 2D class averages of Nb105:RBD:Nb21 complex. (D) Gold-standard Fourier shell correlation (FSC) and Euler angular distribution. (E) Local resolution estimation for Nb105:RBD:Nb21 complex. (F) Rigid docking of Nb105:RBD complex to the interface of the dimeric S. The interface highlighted with the green line is between the Nb framework and RBS.

**Figure S9.**
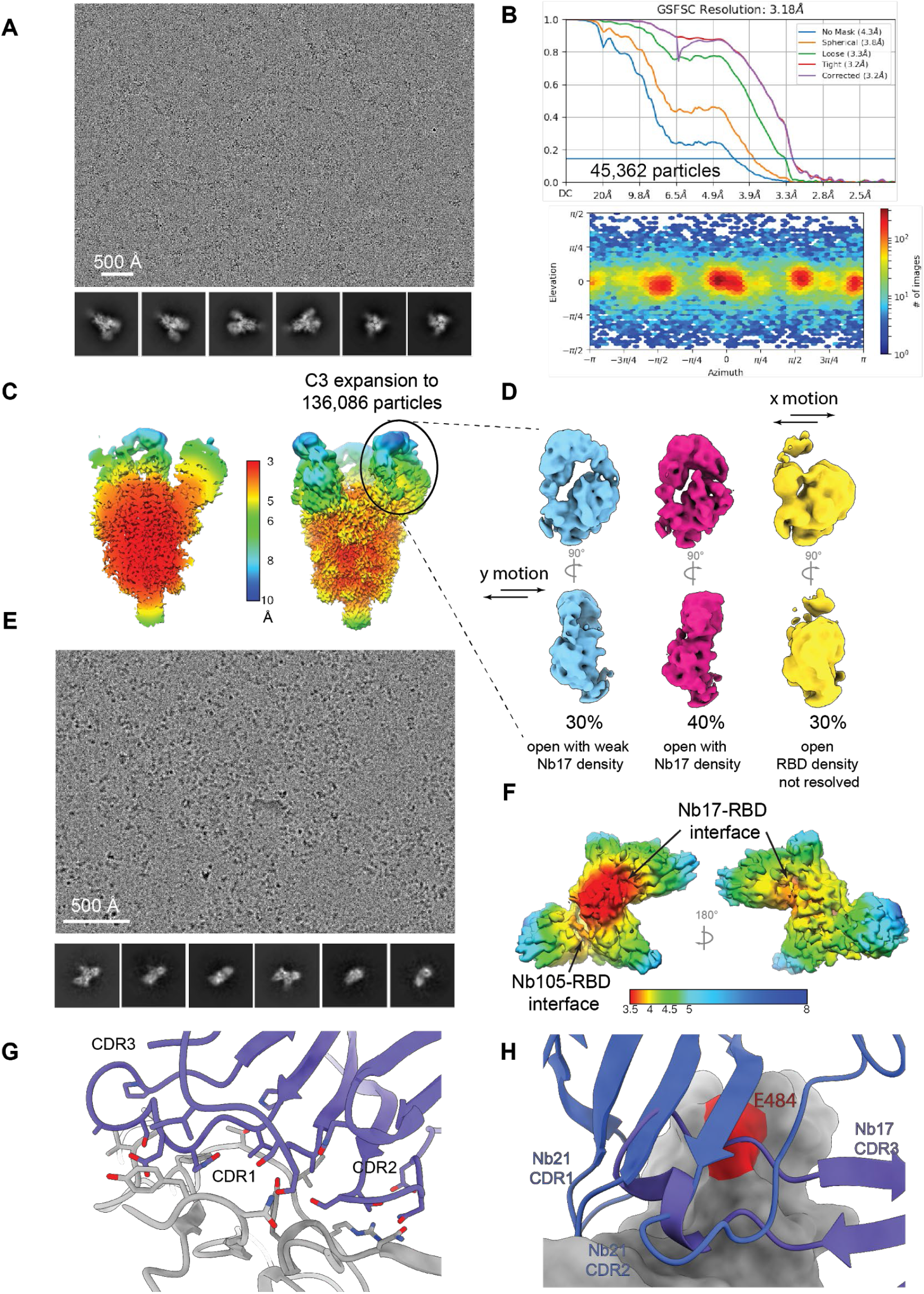
Cryo-EM analysis of Nb17:S and Nb17:RBD:Nb105 complexes (related to Figure 4). (A) Representative micrograph and 2D class averages of Nb17:S complex. (B) Gold-standard Fourier shell correlation (FSC) and Euler angular distribution. (C) Local resolution estimation for Nb17:S complex. (D) Focused classification of the flexible region in Nb17:S complex. The density of Nb17 in class 1 (cyan) is smeared due to motion along the y-direction, class 2 (magenta) has well resolved RBD, Nb17, and NTD density, and both densities of RBD and Nb17 is lost due to motion along the x-direction. (E) Representative micrograph and 2D class averages of Nb17:RBD: Nb105 sample. (F) Local resolution estimation for Nb105:RBD: Nb21 sample. (G) Interface residues of Nb17:RBD complex. (H) Alignment of Nb17:RBD to Nb21:RBD showing the large overlap between Nb17 CDR3 with Nb21 CDR2 and partially Nb21 CDR1.

**Figure S10.**
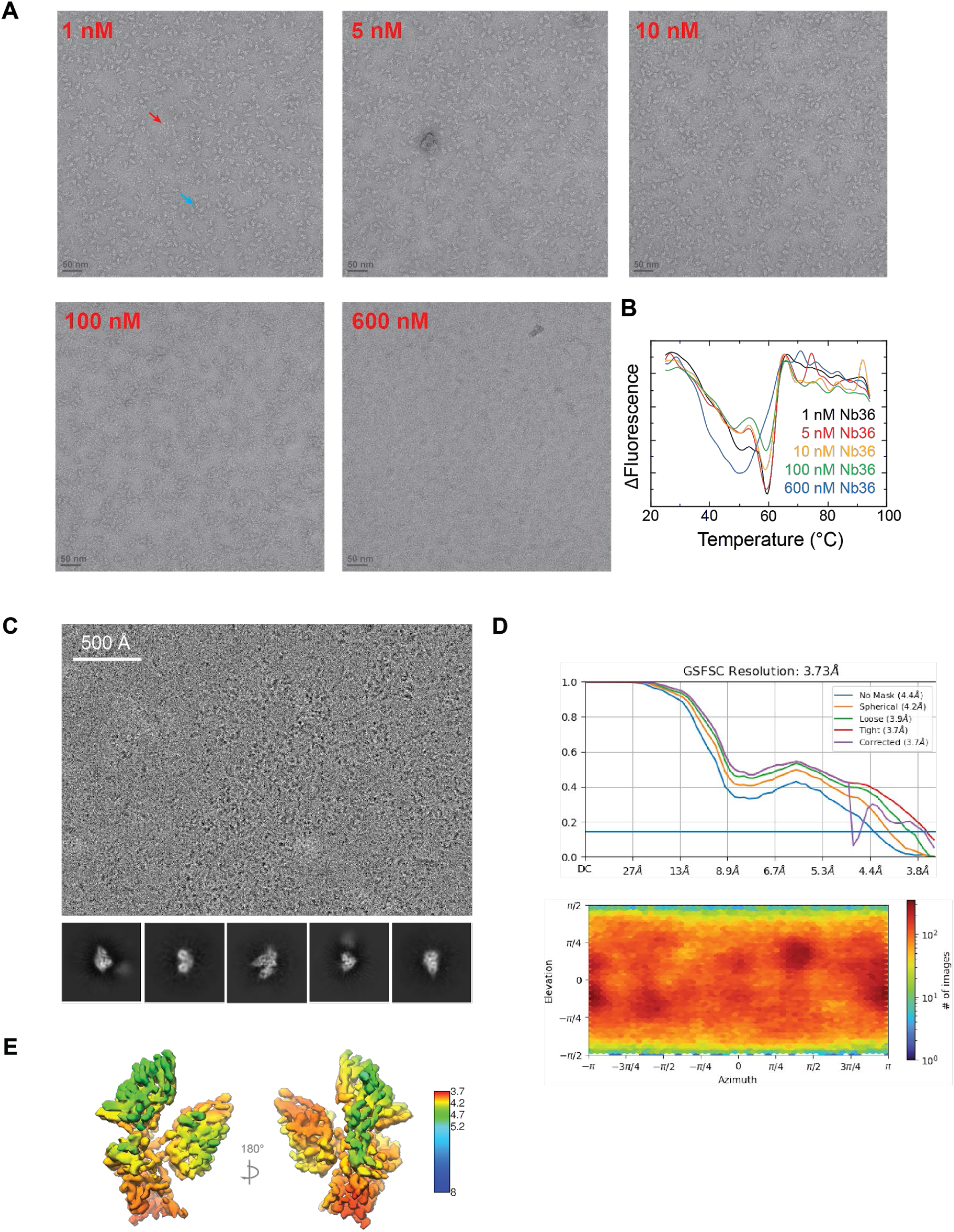
EM Analysis of Nb36 with S and RBD (related to Figure 4). (A) Representative negative stain EM micrographs of spike protein in the presence of an increased concentration of Nb36. An example of an intact trimeric spike particle is highlighted by a blue arrow, and an example of a disrupted spike particle is highlighted by a red arrow. (B) Thermal melting profile of S protein in the presence of an increased concentration of Nb36. (C) Representative micrograph and 2D class averages of Nb36:RBD: Nb21 complex. (D) Gold-standard Fourier shell correlation (FSC) and Euler angular distribution. (E) Local resolution estimation for Nb36:RBD: Nb21 complex.

**Figure S11.**
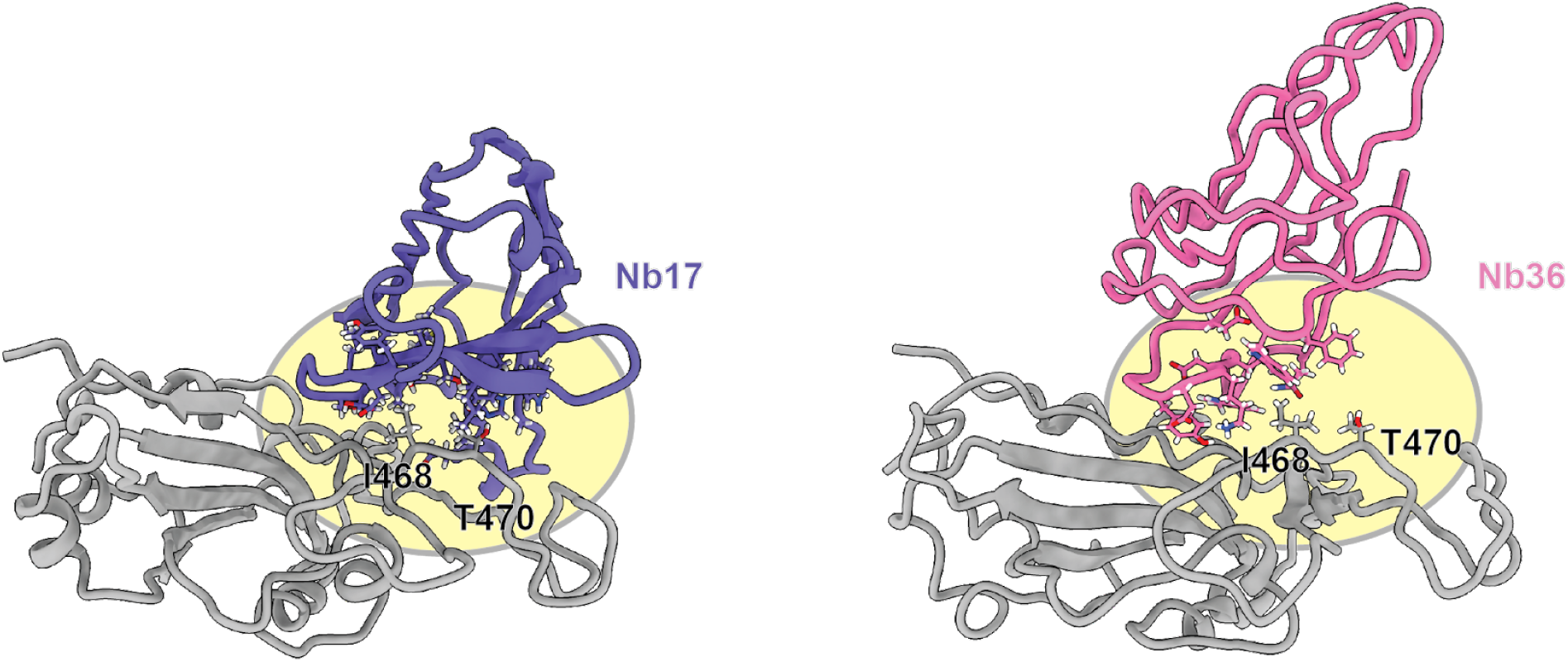
Analysis of the interactions between the RBD residues of I468 and T470 and highly potent neutralizing Nbs from class III (related to Figure 4).

**Figure S12.**
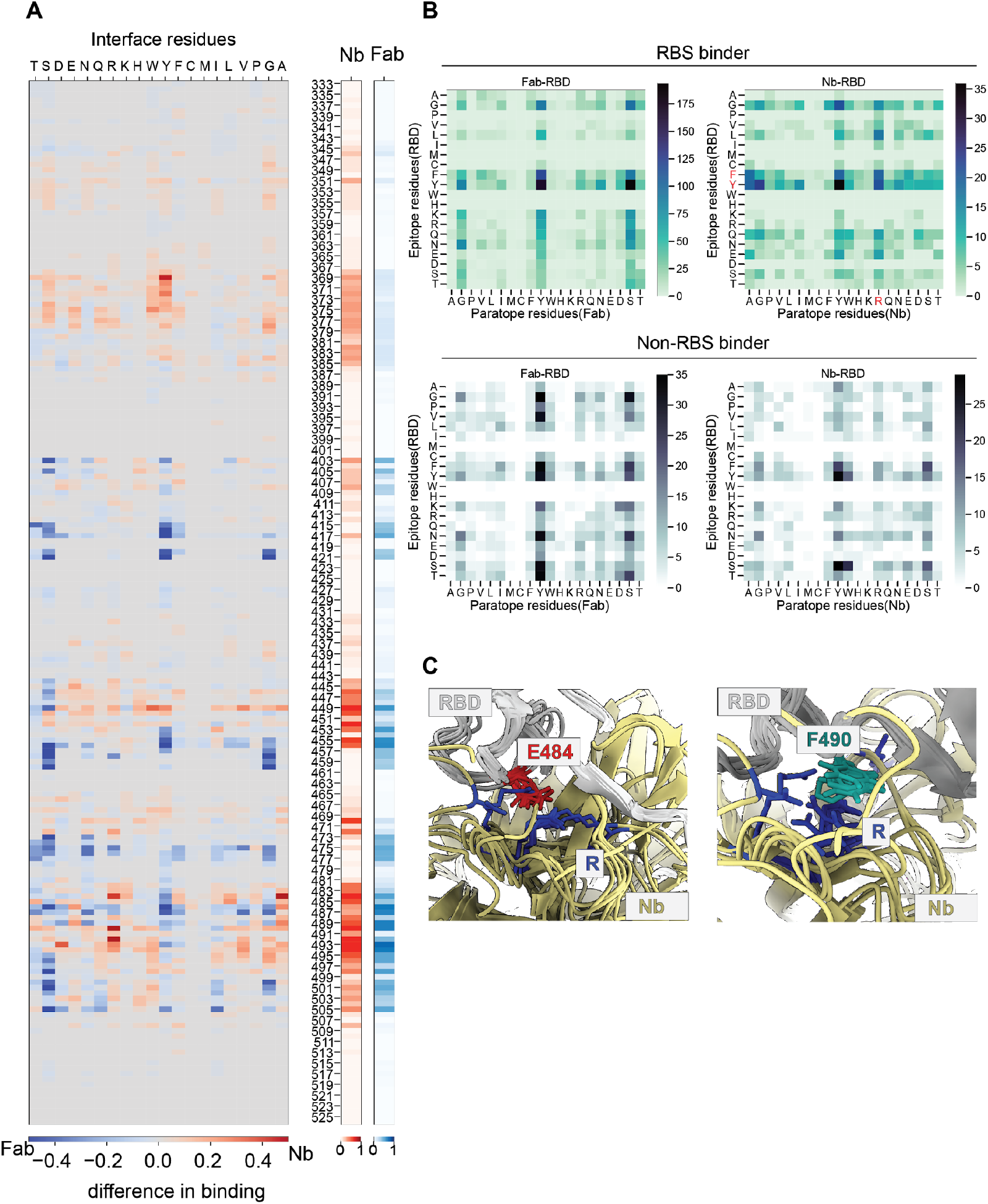
Comparison of neutralizing Nbs and mAbs for RBD binding (related to Figure 7). (A) The heatmap shows the binding difference between Nbs and Fabs in terms of paratope residue utility despite overall similar epitope regions. (B) Heatmaps show the difference in preference of epitope-paratope residues between Nbs and Fabs. The comparisons were made separately for RBS binders and non-RBS binders. Nbs with at least 30% overlapping residues with ACE2 binding sites were considered RBS binders. (C) Illustrations of dominated electrostatic interactions formed between arginine from Nb CDRs and RBD residues. RBD was colored in dark gray, Nbs were colored in khaki, E484 (RBD) was colored in red, F490 (RBD) was colored in teal and R (Nb CDRs) was colored in blue.

**Figure S13.**
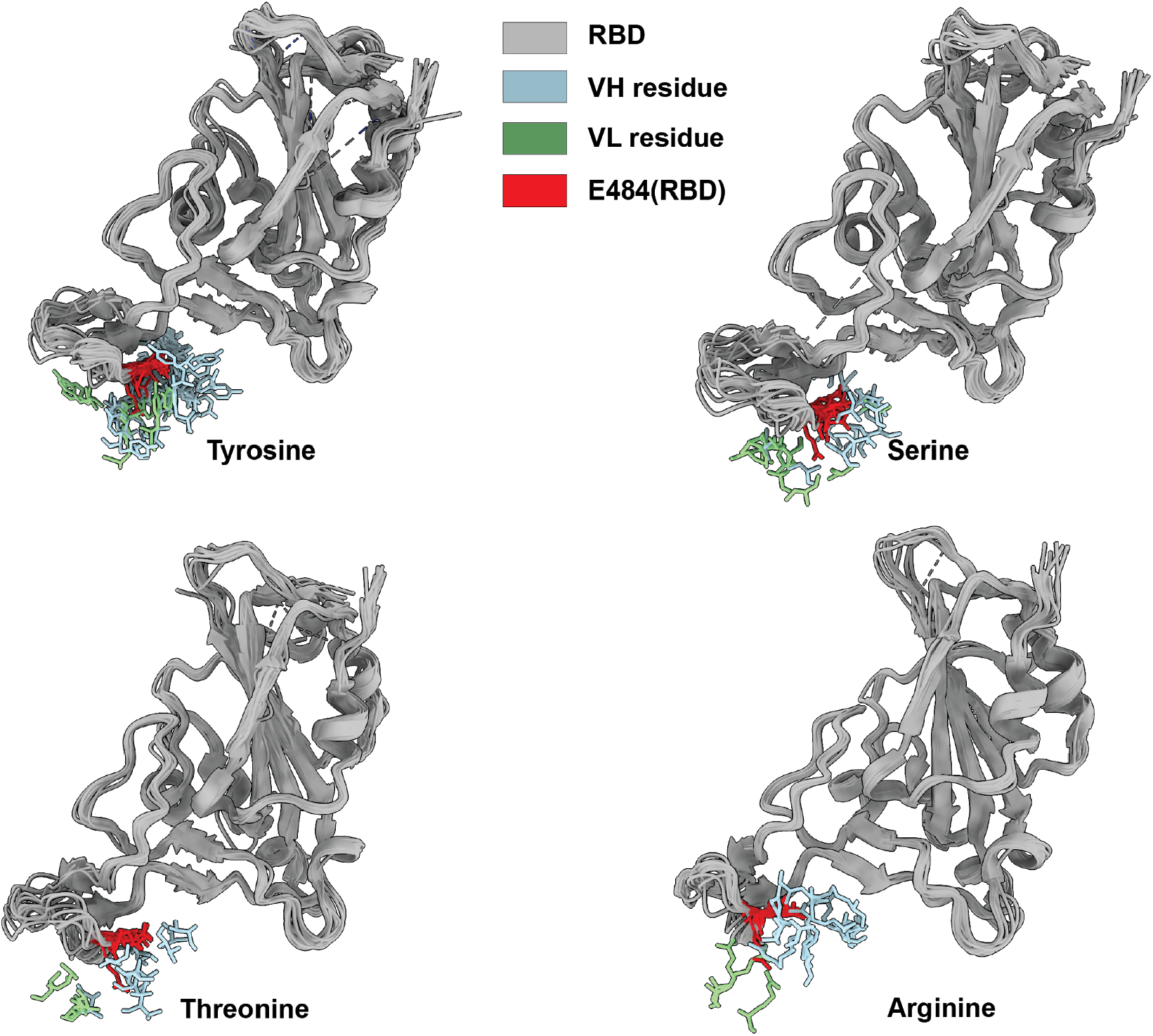
Analysis of interactions of E484 (RBD) with neutralizing Nbs and mAbs (related to Figure 7). Superposition of Fab-RBD structures showing E484 (RBD) forms hydrogen and/or hydrophobic interactions with the respective residues of Fabs. The side chains of residues tyrosine, serine, threonine and arginine in close contact E484 are shown in stick representation. RBD: dark gray, Fab VH: light blue, Fab LH: light green, and residue E484 (RBD): red.

## Tables S1-4

**Table S1. Summary of all RBD mutants, related to Figure 1.**

**Table S2. Statistics for 3D reconstruction and model refinement for Nb:S complexes, related to Figures 2, 3 and 4.**

**Table S3. Statistics for 3D reconstruction and model refinement for 2Nbs:RBD complexes, related to Figures 3 and 4.**

**Table S4. Summary of structural comparisons between mAbs and Nbs, related to Figure 6 and 7.**

## Methods

### Protein expression and purification

The plasmid with cDNA encoding SARS-Cov-2 spike HexaPro (S) was obtained from Addgene. To express the S protein, HEK293-ES cells were transiently transfected with the plasmid using polyethyleneimine and 3.5 mM valproic acid sodium salt to enhance protein production. After 3 hours of transfection, 1 μM kifunensine was added to further boost protein expression. Cell culture was harvested three days after transfection and the supernatant was collected by high-speed centrifugation at 13,000 rpm for 30 mins. The secreted S protein in the supernatant was purified using Ni-NTA agarose columns. Protein eluates were then concentrated and further purified by size-exclusion chromatography using a Superose 6 10/300 column (Cytiva) in a buffer composed of 20 mM Hepes pH 7.5 and 200 mM NaCl. The purified S protein was then pooled and concentrated to 1mg/ml.

The receptor-binding domain (RBD) was expressed and purified as described previously^8^. Briefly, RBD was expressed in Sf9 insect cells as a secreted protein using the baculovirus method. A FLAG-tag and an 8x His-tag were fused to the N terminus of the RBD sequence, and a TEV protease cleavage site was inserted between the His-tag and RBD. The protein was purified by nickel-affinity resins, followed by overnight TEV protease treatment and size exclusion chromatography (Superdex 75). The purified protein was concentrated in a buffer containing 20 mM Hepes pH 7.5 and 150 mM NaCl.

Nanobody genes were codon-optimized and synthesized by Synbio as previously described^8^. All nanobody sequences were cloned into pET-21b(+) vectors using EcoRI and HindIII restriction sites. Plasmids were transformed into BL21 (DE3) cells and plated onto Agar gel media with 50 μg/ml ampicillin. Agar plates were incubated at 37°C overnight, and single colonies were picked for protein purification. The cell culture was allowed to grow at 37°C to an OD600 of 0.5-0.6, at which point the temperature was lowered to 16°C and 0.5-1mM IPTG was added to induce protein expression overnight. Cells were then pelleted, resuspended in a lysis buffer (1x PBS, 150 mM NaCl, 0.2% Triton-X100, and protease inhibitors), and ultrasonicated on ice. The clarified cell lysate was collected by centrifugation at 15,000 x g for 10 mins. His-tagged nanobodies were captured using cobalt resin and eluted with a pre-chilled buffer containing imidazole (50 mM NaPO4, 300 mM NaCl, 150 mM Imidazole, pH 7.4). Nanobodies eluted from His-Cobalt resin were further purified using a Superdex 75 gel filtration column using filtered 1x PBS. Nanobodies were used fresh or flash frozen and stored at −80°C before use.

### Cryo-EM sample preparation and imaging

For S complexed with Nb21, Nb34, and Nb95, each Nb was mixed with the S protein at a molecular ratio of 5 to 1 and incubated at 4 degrees for 30 minutes. The complex was then diluted in a buffer containing 20 mM Hepes pH 7.5 and 200 mM NaCl to reach a concentration of the S protein at 0.2 mg/ml. Then the sample was applied to a 1.2/1.3 UltrAuFoil grid (Electron Microscopy Sciences) that had been freshly glow-discharged and plunge-frozen in liquid ethane using an FEI Vitrobot Mark IV. All cryo-EM data were collected on Titan Krios transmission electron microscopes (Thermo Fisher) operating at 300 kV. For the S and Nb21 complex, images were acquired on a Falcon 3 detector, with a nominal magnification of 96,000, corresponding to a final pixel size of 0.83 Å/pixel. For each image stack, a total dose of about 62 electrons was equally fractionated into 70 fractions with ~0.88 e^-^/Å2/fraction. EPU 2 Software was used to automate data collection. Defocus values used to collect the dataset ranged from −0.5 to −3.5 μm.

For the complexes of S with Nb95 and Nb34, sample preparation and data collection were similar to those for the complex of S with Nb21 except that the data were acquired on a Gatan K3-Summit detector. Further details of data collection parameters are summarized in **Table S2 and S3.**

For S complexes with Nb17 and 105, purified nanobodies were mixed with the SARS-CoV-2 S HexaPro trimer with 2:1 molar ratio nanobodies to a final concentration of 0.1 mg/mL S protein and incubated at room temperature for two hours. Cryo-EM grids (Quantifoil AU 1.2/1.3 300 mesh) were glow-discharged and coated with graphene oxide thin layer flakes following the protocol from reference^43^ (figshare. Media. https://doi.org/10.6084/m9.figshare.3178669.v1). The cryo-EM specimens were prepared using an FEI Vitrobot Mark IV with 3.5μl of freshly prepared nanobody:S complex. Grids were blotted for 3 s with blot force −5 in 100% humidity at 4°C prior to plunge freezing. The frozen-dehydrated grids were transferred to a Titan Krios (Thermo Fisher Scientific) transmission electron microscope equipped with a Gatan K3direct-electron counting camera and BioQuantum energy filter for data acquisition. Movies of the specimen were recorded with a nominal defocus setting in the range of −0.5 to −2.0 μm using SerialEM with beam-tilt image-shift data collection strategy with a 3 x 3 pattern and 1 shot per hole. The movie stacks were collected in the correlated double sampling (CDS) super-resolution mode of the K3 camera at a nominal magnification of 81,000 yielding a physical pixel size of 1.08 Å/pixel. Each stack was exposed for 5 s, with each frame exposed for 0.1 s, resulting in a 50-frame movie. For datasets without using CDS mode, the movie stacks were collected in the super-resolution mode at a nominal magnification of 81,000 with an exposure time of 2.5 s, and each frame exposed for 0.05 s. The total accumulated dose on the specimen was 40 e/Å^2^ for each stack.

For the trimeric Nb complexes (Nb105:RBD: Nb21, Nb17:RBD: Nb105 and Nb36:RBD: Nb21), two purified Nbs were mixed with purified RBD with 1.1:1.1:1 molar ratio and subsequently polished by size-exclusion chromatography (SEC). Peak fraction corresponding to the trimeric complexes was used for cryo-grid preparation. Movies of the specimen were recorded with a nominal defocus setting in the range of −0.5 to −2.5 μm using SerialEM with beam-tilt image-shift data collection strategy with a 3 x 3 pattern and 3 shot per hole. The movie stacks were collected in the correlated double sampling (CDS) super-resolution mode of the K3 camera at a nominal magnification of 165,000 yielding a physical pixel size of 0,52 Å/pixel. Each stack was exposed for 2.8 s, with each frame exposed for 0.1 s, resulting in a 28-frame movie. The total accumulated dose on the specimen was 108 e/Å^2^ for each stack.

### Cryo-EM data processing

For the samples of S protein with Nb21, Nb34, and Nb95, the cryo-EM data processing was performed using Relion 3.1. Beam-induced motion correction was performed using the motion correction program implemented in Relion to generate average micrographs and dose-weighted micrographs from all frames. Contrast transfer function (CTF) parameters were estimated using CTFFIND4 from average micrographs. The loG-based auto-picking procedure was used for reference-free particle picking. Initial particle stacks were subjected to 2D classification and the best class averages that represented different views were selected as templates for second round automatic particle picking from the dose-weighted micrographs.

For the S protein with Nb21 data, approximately 900,000 particles were auto-picked from 2,574 micrographs for further processing. The whole set of particles was cleaned to remove contaminants or junk particles by 2D classification and 3D classification using 2x binned particles. Finally, approximately 135,000 particles were used for 3D auto-refinement with the structure of EMD-22221 (EMBD ID) low-pass filtered to 40Å as the reference. This yielded a map of ~3.4 Å resolution (corrected gold-standard FSC 0.143 criterion). The particles were re-extracted and used for further 3D classification into four classes. The most populated two classes, which contained 53% (1-up-down RBDs) and 29% (2-up-1-down RBDs) of particles were subjected to further 3D auto-refinement. To improve the local density of Nb21 and RBD, focused refinement was performed with a soft mask applied to one down RBD and Nb21, resulting in an improved local resolution ranging from 4.5 Å to 3.3 Å for the RBD and Nb21 interface. All maps were sharpened using the post-processing program in Relion or DeepEMhancer. The local resolution was estimated by ResMap in Relion. Similar approaches were used to solve the structures of S protein with Nb95 and Nb34 as well. The detailed information of data processing is shown in **Figures S4 and S6–10** and **Table S2-3**.

For other structures, each movie stack was processed on-the-fly using CryoSPARC live (version 3.0.0) ^44,45^. The movie stacks were aligned using patch motion correction with an F-crop factor of 0.5. The contrast-transfer function (CTF) parameters of each particle were estimated using patch CTF. Particles were auto-picked using a 220 and 100 Å gaussian blob for Nb:S and 2Nbs: RBD complexes respectively. The numbers of bin2 particles selected after 2D classification are included in **Table S2**. The initial 3D volume and decoys were generated using *ab initio* reconstruction with a minibatch size of 1000 using a set of rebalanced 2D classes. The particles after 2D clean-up were submitted to one round of heterogeneous refinement with *ab initio* 3D volume from good 2D classes and decoy 3D volumes from bad 2D classes. Based on the coordinates and angular information of these particles, bin1 particles of the 3D class with well-resolved secondary structure features were re-extracted from the dose-weighted micrographs. For small trimeric complexes, a pixel size that can achieve the resolution limit of the sample, instead of bin1 pixel, was used for the final reconstruction to prevent overfitting. The final particle set was subjected to non-uniform 3D refinements^45^, followed by local 3D refinements, yielding final maps with reported global resolutions using the 0.143 criteria of the gold-standard Fourier shell correlation (FSC) (**Table S2**). The half maps were used to determine the local resolution of each map and focused classification was performed using Relion 3.0 ^46,47^. For Nb17:S complex, the final particles (45,362) were aligned to the C3 symmetry axis to expand the particle set to 136,086 (**Figure S9C**). Then a mask focused on the arc shape including RBD, Nb17 and NTD was created with the binary map extended 10 pixels and a soft-edge of 10 pixels. The cryosparc particle set was converted to relion star file using pyem (https://github.com/asarnow/pyem), and focused classification was performed by Relion with k=3. The class with densities for all three targeted domains well resolved was selected for further local refinement in CryoSPARC.

### Model building and structure refinement

For modeling whole S protein with Nbs, the RBD models were generated by docking the atomic model of SARS-Cov2 RBD (PDB ID 7JVB, chain B) into the refined cryo-EM density using Chimera (UCSF). Nb structures were modeled ab initio in Coot using based on the locally refined cryo-EM maps and refined in Phenix. After refinement, each residue of the sequence-updated models was manually checked and refined iteratively in Coot and Phenix. Structural models were validated by MolProbity. The final refinement statistics are listed in **Table S2-3**.

### CDR3 loop modeling

To optimize the CDR3 loop conformation, it was modeled *ab-initio* using ‘RosettaAntibody3’ H3 loop modeling with restraints (distance, dihedral, and planar angles ^48^) generated by NanoNet. NanoNet is a deep residual neural network, similar to DeepH3 ^49^, trained on solved CDR3 loops of antibodies and nanobodies from the PDB. NanoNet uniqueness comes from the fact that it takes as input only the sequence of the CDRs (each in a single one-hot encoding matrix) without the framework region. In addition, it uses MSE loss and predicts the pairwise distances and angles directly (for angles, it predicts the *sine* and *cosine* values to overcome cyclic loss), instead of using categorical cross-entropy loss and trying to predict the pairwise probability distributions. NanoNet architecture consists of two 2D residual blocks, followed by two convolutional layers for each output, with the *tanh* activation function for angles and ReLU (rectified linear activation function) for distances.

For each nanobody, 100 models were generated, and the one that fitted best in the cryo-EM density map was chosen manually. For Nb20, Nb21, Nb95, Nb105, Nb34 the models generated were similar to the ones without the optimization. For Nbs 17 and 36, the models generated from ‘RosettaAntibody3’ with NanoNet fitted better in the cryo-EM map and were further refined in the density map.

### Contact heat map

An RBD residue and an Ab/Nb residue were defined in contact if the distance between any pair of their atoms was lower than a threshold of 6 Å. The Ab/Nb contact value of each RBD residue is calculated as the average of all the Ab/Nb contacts. Nb classes were clustered using k-means (k=3). The conservation score was obtained from Consurf by querying the RBD sequence.

### Measurement of buried surface area(BSA)

The solvent-accessible surface area(SASA) of molecules was calculated by FreeSASA^50^. The buried surface area in the case of the Nb-RBD complex was then calculated by *BSA* = 1/2[*SASA*(*Nb*) + *SASA*(*RBD*) – *SASA* (*complex*)].

### Measurement of structural overlap between Nb and corresponding best matched Fab

The best matched Fab for an Nb was obtained using the epitope similarity(Jaccard-index). The Nb-RBD complex structure was superimposed on its best matched Fab-RBD structure and protein volume was calculated using ProteinVolume^51^. Then the structural overlap was calculated by *structural overlap* = [*Volume*(*Nb*) + *Volume*(*RBD*) — *Volume*(*complex*)]/*Volume*(*RBD*).

### Measurement of the interface curvature

The interface curvature was calculated as the average of the shape function of the interface atoms of the antigen or the Nb. For this purpose, a sphere of radius R (6Å) is placed at a surface point of the interface atom. The fraction of the sphere inside the solvent-excluded volume of the protein is the shape function at the corresponding atom ^52^.

### ELISA (Enzyme-Linked Immunosorbent Assay)

Antigens (RBD and RBD variants) were coated onto 96-well ELISA plates, with 150 ng of protein per well in the coating buffer (15 mM sodium carbonate, 35mM Sodium Bicarbonate, pH 9.6) at 4°C for overnight. The plates were decanted, washed with a buffer (1x PBS, 0.05% Tween 20), and blocked for 2 hours at room temperature (1x PBS, 0.05% Tween 20, 5% milk powder). Nanobodies were serially 5x diluted in blocking buffers from 500 nM to 6.4 pM. Anti-T7 tag HRP-conjugated secondary antibodies were diluted at 1:5000 and incubated at room temperature for 1 hour. Upon washing, samples were further incubated in the dark for 10 minutes with freshly prepared 3,3’,5,5’-Tetramethylbenzidine (TMB) substrate. Upon quenching the reaction with a STOP solution, the plates were measured at wavelengths of 450 nm with background subtraction at 550 nm. The raw data was processed and fitted into the 4PL curve using the Prism Graphpad 9.0. IC50s were calculated and fold changes of binding affinity were calculated to generate the heatmap.

For ACE2 competitive ELISA assays, the super stable spike was coated at 80 ng/ml on the plate. Nanobodies were serially 5x diluted in blocking buffers from 500 nM to 32 pM with an addition of 60 ng/well of biotinylated hACE2 for competition. No Nb was used as a negative control. Pierce High Sensitivity Neutravidin-HRP antibodies were used at 1:8000. The hACE2 percentage was calculated by the reading at each Nb concentration divided by the reading at the negative control. Then the data processed and fitted into the 4PL curve using the Prism Graphpad 9.0.

### Pseudovirus neutralization assay

The 293T-hsACE2 stable cell line and the pseudotyped SARS-CoV-2 particles (wild-type and mutants) with luciferase reporters were purchased from the Integral Molecular. The B.1.1.7 UK pseudotyped virus contains all of the naturally prevalent mutations for that strain. The SA 501Y.V2 contains all of the naturally prevalent mutations except del241-243, which is replaced by an L242H substitution for the pseudovirus (**Figure S2**). The neutralization assay was carried out according to the manufacturers’ protocols in duplicates. In brief, 3-fold or 5-fold serially diluted Nbs were incubated with the pseudotyped SARS-CoV-2-luciferase for 1 hour at 37 °C. At least eight concentrations were tested for each Nb. Pseudovirus in culture media without Nbs was used as a negative control. 100 μl of the mixtures were then incubated with 100 μl 293T-hsACE2 cells at 3×10e5 cells/ml in the 96-well plates. The infection took ~72 hours at 37 °C with 5% CO2. The luciferase signal was measured using the *Renilla*-Glo luciferase assay system with the luminometer at 1 ms integration time. The obtained relative luminescence signals (RLU) from the wells were normalized according to the negative control and the neutralization percentage was calculated at each concentration. The data was then processed by Prism GraphPad 9.0 to fit into a 4PL curve and to calculate the logIC50 (half-maximal inhibitory concentration).

### Protein thermal shift assay

Thermal denaturation of S protein in the presence of an increased concentration of Nb36 was monitored by differential scanning fluorimetry using Protein Thermal Shift^™^ dye kit^53^. Briefly, the same protein samples used for negative stain EM were diluted to a final assay concentration of 100 nM in PBS with 1 mM DTT and 1:1000 fluorescence dye (TFS 4461146). The final assay volume was 20 μL, with 1, 5, 10, 100, and 600 nM of Nb36 was added to a final concentration of 100 nM S protein. Heat denaturation curves were recorded using a realtime PCR instrument (StepOne^™^) applying a temperature gradient of 1 °C/min. Analysis of the data was performed using Excel. Melting temperatures of protein samples were determined by the inflection points of the plots of –d(RFU)/dT.

### Negative-stain electron microscopy

For negative staining electron microscopy, 3 μl of specified concentration of Nb36 with the S protein was applied to a glow-discharged grid coated with carbon film. The sample was left on the carbon film for 60s, followed by negative staining with 2% uranyl formate. Electron microscopy micrographs were recorded on a Gatan Ultrascan CCD camera at 22,000 × magnification in an FEI Tecnai 12 electron microscope operated at 100 keV.

## Acknowledgments

We are grateful to the Cryo-Electron Microscopy Core at the CWRU School of Medicine and K. Li for access to the sample preparation and cryo-EM instrumentation. Computational support was provided by the Case Western Reserve University High-Performance Computing Cluster. This research was supported in part by the National cryo-EM Facility of the National Cancer Institute at the Frederick National Laboratory for Cancer Research under contract HSSN261200800001E. This work was performed in part at the National Center for cryo-EM Access and Training (NCCAT) and the Simons Electron Microscopy Center located at the New York Structural Biology Center, supported in part by the NIH Common Fund Transformative High-Resolution Cryo-Electron Microscopy program (U24 GM129539,) and by grants from the Simons Foundation (SF349247) and NY State Assembly. This work was further supported by grants from the NIH (R35 GM137905 to Y.S., R35 GM128641 to C.Z., R01 GM133841, and R01 CA240993 to D.J.T.), a CTSI grant (Y.S.), and ISF 1466/18, Israel ministry of Science and Technology and HUJI-CIDR (D.S.). We thank Gideon Schreiber (Weizmann Institute of Science) for providing B62 and Alex Guseman to help.

## Author contributions

Y.S. conceived the study. W.H., D.S., C.Z., J.S., D.T., A.K. B.A.H, J.S. and J.C collected and analyzed the cryo-EM data. W.H., Y.J.K, and Y.X. prepared proteins and performed biochemical and functional assays. Z.S., D.S., Y.S., T.C., and W.H. systematically analyzed and compared antibody structures. W.H., Z.S., D.S., C.Z., Y.S., prepare figures. Y.S. and W.H. wrote the manuscript with substantial input from D.S. and C.Z. All authors edited the manuscript.

## Notes

### Competing Interest Statement

University of Pittsburgh has filed a patent application related to the nanobodies of this study

## References

1 Huang, A. T. et al. A systematic review of antibody mediated immunity to coronaviruses: kinetics, correlates of protection, and association with severity. Nat Commun 11, 4704, doi:10.1038/s41467-020-18450-4 (2020).

2 Cohen, M. S. Monoclonal Antibodies to Disrupt Progression of Early Covid-19 Infection. N Engl J Med 384, 289–291, doi:10.1056/NEJMe2034495 (2021).

3 Chen, P. et al. SARS-CoV-2 Neutralizing Antibody LY-CoV555 in Outpatients with Covid-19. N Engl J Med 384, 229–237, doi:10.1056/NEJMoa2029849 (2021).

4 Weinreich, D. M. et al. REGN-COV2, a Neutralizing Antibody Cocktail, in Outpatients with Covid-19. N Engl J Med 384, 238–251, doi:10.1056/NEJMoa2035002 (2021).

5 Koenig, P. A. et al. Structure-guided multivalent nanobodies block SARS-CoV-2 infection and suppress mutational escape. Science 371, doi:10.1126/science.abe6230 (2021).

6 Bracken, C. J. et al. Bi-paratopic and multivalent VH domains block ACE2 binding and neutralize SARS-CoV-2. Nat Chem Biol 17, 113–121, doi:10.1038/s41589-020-00679-1 (2021).

7 Schoof, M. et al. An ultrapotent synthetic nanobody neutralizes SARS-CoV-2 by stabilizing inactive Spike. Science 370, 1473–1479, doi:10.1126/science.abe3255 (2020).

8 Xiang, Y. et al. Versatile, Multivalent Nanobody Cocktails Efficiently Neutralize SARS-CoV-2. bioRxiv, doi:10.1101/2020.08.24.264333 (2020).

9 Walter, J. D. et al. Highly potent bispecific sybodies neutralize SARS-CoV-2. bioRxiv, 2020.2011.2010.376822, doi:10.1101/2020.11.10.376822 (2020).

10 Ahmad, J., Jiang, J., Boyd, L. F., Natarajan, K. & Margulies, D. H. Synthetic nanobody-SARS-CoV-2 receptor-binding domain structures identify distinct epitopes. bioRxiv, doi:10.1101/2021.01.27.428466 (2021).

11 Custodio, T. F. et al. Selection, biophysical and structural analysis of synthetic nanobodies that effectively neutralize SARS-CoV-2. Nat Commun 11, 5588, doi:10.1038/s41467-020-19204-y (2020).

12 Huo, J. et al. Neutralizing nanobodies bind SARS-CoV-2 spike RBD and block interaction with ACE2. Nat Struct Mol Biol 27, 846–854, doi:10.1038/s41594-020-0469-6 (2020).

13 Lu, Q. et al. Development of multivalent nanobodies blocking SARS-CoV-2 infection by targeting RBD of spike protein. J Nanobiotechnology 19, 33, doi:10.1186/s12951-021-00768-w (2021).

14 Nambulli, S. et al. Inhalable Nanobody (PiN-21) prevents and treats SARS-CoV-2 infections in Syrian hamsters at ultra-low doses. bioRxiv, doi:10.1101/2021.02.23.432569 (2021).

15 Bangaru, S. et al. Structural analysis of full-length SARS-CoV-2 spike protein from an advanced vaccine candidate. Science 370, 1089–1094, doi:10.1126/science.abe1502 (2020).

16 Casalino, L. et al. AI-Driven Multiscale Simulations Illuminate Mechanisms of SARS-CoV-2 Spike Dynamics. bioRxiv, doi:10.1101/2020.11.19.390187 (2020).

17 Lu, M. et al. Real-Time Conformational Dynamics of SARS-CoV-2 Spikes on Virus Particles. Cell Host Microbe 28, 880–891 e888, doi:10.1016/j.chom.2020.11.001 (2020).

18 Cai, Y. et al. Distinct conformational states of SARS-CoV-2 spike protein. Science 369, 1586–1592, doi:10.1126/science.abd4251 (2020).

19 Wang, P. et al. Increased Resistance of SARS-CoV-2 Variants B.1.351 and B.1.1.7 to Antibody Neutralization. bioRxiv, doi:10.1101/2021.01.25.428137 (2021).

20 Wang, Z. et al. mRNA vaccine-elicited antibodies to SARS-CoV-2 and circulating variants. Nature, doi:10.1038/s41586-021-03324-6 (2021).

21 Zhou, D. et al. Robust SARS-CoV-2 infection in nasal turbinates after treatment with systemic neutralizing antibodies. Cell Host Microbe, doi:10.1016/j.chom.2021.02.019 (2021).

22 Davies, N. G. et al. Estimated transmissibility and impact of SARS-CoV-2 lineage B.1.1.7 in England. Science, doi:10.1126/science.abg3055 (2021).

23 Wibmer, C. K. et al. SARS-CoV-2 501Y.V2 escapes neutralization by South African COVID-19 donor plasma. bioRxiv, doi:10.1101/2021.01.18.427166 (2021).

24 Cele, S. et al. Escape of SARS-CoV-2 501Y.V2 variants from neutralization by convalescent plasma. medRxiv, 2021.2001.2026.21250224, doi:10.1101/2021.01.26.21250224 (2021).

25 Thomson, E. C. et al. Circulating SARS-CoV-2 spike N439K variants maintain fitness while evading antibody-mediated immunity. Cell, doi:10.1016/j.cell.2021.01.037 (2021).

26 Xiang, Y. et al. Integrative proteomics identifies thousands of distinct, multi-epitope, and high-affinity nanobodies. Cell Syst, doi:10.1016/j.cels.2021.01.003 (2021).

27 Zahradník, J. et al. SARS-CoV-2 RBD in vitro evolution follows contagious mutation spread, yet generates an able infection inhibitor. bioRxiv, 2021.2001.2006.425392, doi:10.1101/2021.01.06.425392 (2021).

28 Hsieh, C. L. et al. Structure-based design of prefusion-stabilized SARS-CoV-2 spikes. Science 369, 1501–1505, doi:10.1126/science.abd0826 (2020).

29 Barnes, C. O. et al. Structures of Human Antibodies Bound to SARS-CoV-2 Spike Reveal Common Epitopes and Recurrent Features of Antibodies. Cell 182, 828–842 e816, doi:10.1016/j.cell.2020.06.025 (2020).

30 Yuan, M. et al. A highly conserved cryptic epitope in the receptor binding domains of SARS-CoV-2 and SARS-CoV. Science 368, 630–633, doi:10.1126/science.abb7269 (2020).

31 Huo, J. et al. Neutralization of SARS-CoV-2 by Destruction of the Prefusion Spike. Cell Host Microbe 28, 445–454 e446, doi:10.1016/j.chom.2020.06.010 (2020).

32 Starr, T. N. et al. Deep Mutational Scanning of SARS-CoV-2 Receptor Binding Domain Reveals Constraints on Folding and ACE2 Binding. Cell 182, 1295–1310 e1220, doi:10.1016/j.cell.2020.08.012 (2020).

33 Greaney, A. J. et al. Complete Mapping of Mutations to the SARS-CoV-2 Spike Receptor-Binding Domain that Escape Antibody Recognition. Cell Host Microbe 29, 44–57 e49, doi:10.1016/j.chom.2020.11.007 (2021).

34 Greaney, A. J. et al. Comprehensive mapping of mutations in the SARS-CoV-2 receptor-binding domain that affect recognition by polyclonal human plasma antibodies. Cell Host Microbe, doi:10.1016/j.chom.2021.02.003 (2021).

35 Mitchell, L. S. & Colwell, L. J. Analysis of nanobody paratopes reveals greater diversity than classical antibodies. Protein Eng Des Sel 31, 267–275, doi:10.1093/protein/gzy017 (2018).

36 Sztain, T. et al. A glycan gate controls opening of the SARS-CoV-2 spike protein. bioRxiv, 2021.2002.2015.431212, doi:10.1101/2021.02.15.431212 (2021).

37 Lucy, F. et al. Free Energy Landscapes for RBD Opening in SARS-CoV-2 Spike Glycoprotein Simulations Suggest Key Interactions and a Potentially Druggable Allosteric Pocket. (2020).

38 Zimmerman, M. I. et al. SARS-CoV-2 Simulations Go Exascale to Capture Spike Opening and Reveal Cryptic Pockets Across the Proteome. bioRxiv, 2020.2006.2027.175430, doi:10.1101/2020.06.27.175430 (2020).

39 Casalino, L. et al. AI-Driven Multiscale Simulations Illuminate Mechanisms of SARS-CoV-2 Spike Dynamics. bioRxiv, 2020.2011.2019.390187, doi:10.1101/2020.11.19.390187 (2020).

40 Chen, Y. et al. Quick COVID-19 Healers Sustain Anti-SARS-CoV-2 Antibody Production. Cell 183, 1496–1507 e1416, doi:10.1016/j.cell.2020.10.051 (2020).

41 Wajnberg, A. et al. Robust neutralizing antibodies to SARS-CoV-2 infection persist for months. Science 370, 1227–1230, doi:10.1126/science.abd7728 (2020).

42 Carfi, A., Bernabei, R., Landi, F. & Gemelli Against, C.-P.-A. C. S. G. Persistent Symptoms in Patients After Acute COVID-19. JAMA 324, 603–605, doi:10.1001/jama.2020.12603 (2020).

43 Bokori-Brown, M. et al. Cryo-EM structure of lysenin pore elucidates membrane insertion by an aerolysin family protein. Nat Commun 7, 11293, doi:10.1038/ncomms11293 (2016).

44 Punjani, A., Rubinstein, J. L., Fleet, D. J. & Brubaker, M. A. cryoSPARC: algorithms for rapid unsupervised cryoEM structure determination. Nat Methods 14, 290–296, doi:10.1038/nmeth.4169 (2017).

45 Punjani, A., Zhang, H. & Fleet, D. J. Non-uniform refinement: adaptive regularization improves single-particle cryo-EM reconstruction. Nat Methods 17, 1214–1221, doi:10.1038/s41592-020-00990-8 (2020).

46 Kimanius, D., Forsberg, B. O., Scheres, S. H. & Lindahl, E. Accelerated cryo-EM structure determination with parallelisation using GPUs in RELION-2. Elife 5, doi:10.7554/eLife.18722 (2016).

47 Zivanov, J. et al. New tools for automated high-resolution cryo-EM structure determination in RELION-3. Elife 7, doi:10.7554/eLife.42166 (2018).

48 Yang, J. et al. Improved protein structure prediction using predicted interresidue orientations. Proc Natl Acad Sci U S A 117, 1496–1503, doi:10.1073/pnas.1914677117 (2020).

49 Ruffolo, J. A., Guerra, C., Mahajan, S. P., Sulam, J. & Gray, J. J. Geometric potentials from deep learning improve prediction of CDR H3 loop structures. Bioinformatics 36, i268–i275, doi:10.1093/bioinformatics/btaa457 (2020).

50 Mitternacht, S. FreeSASA: An open source C library for solvent accessible surface area calculations. F1000Res 5, 189, doi:10.12688/f1000research.7931.1 (2016).

51 Chen, C. R. & Makhatadze, G. I. ProteinVolume: calculating molecular van der Waals and void volumes in proteins. BMC Bioinformatics 16, 101, doi:10.1186/s12859-015-0531-2 (2015).

52 Connolly, M. L. Shape Complementarity at the Hemoglobin Alpha-1-Beta-1-Subunit Interface. Biopolymers 25, 1229–1247, doi:DOI 10.1002/bip.360250705 (1986).

53 Niesen, F. H., Berglund, H. & Vedadi, M. The use of differential scanning fluorimetry to detect ligand interactions that promote protein stability. Nat Protoc 2, 2212–2221, doi:10.1038/nprot.2007.321 (2007).

